# The Holdup Multiplex, an assay for high-throughput measurement of protein-ligand affinity constants using a mass-spectrometry readout

**DOI:** 10.1101/2022.12.08.519103

**Authors:** François Delalande, Gergo Gogl, Aurélien Rohrbacher, Camille Kostmann, Pascal Eberling, Christine Carapito, Gilles Travé, Elodie Monsellier

## Abstract

The accurate description and subsequent modeling of protein interactomes requires quantification of their affinities at proteome-wide scale. Here we develop and validate the Holdup Multiplex, a versatile assay for high-throughput measurement of protein-ligand affinity constants that uses mass-spectrometry as readout. The method can quantify thousands of affinities in one single run, with high precision and over several orders of magnitude. We applied this strategy to the seven human 14-3-3 isoforms, quantifying in a few sample-runs their interaction with 1,000 different phosphopeptides. We were able to identify hundreds of new 14-3-3 binding sites. We showed that the seven human 14-3-3 display similar specificities but staggered affinities, 14-3-3g being always the best binder and 14-3-3ε and σ, the weakest. Finally, we identified dozens of 14-3-3 bindings sites, some intervening in key signaling pathways, that were either stabilized or destabilized by the phytotoxin Fusicoccin-A. Our approach, which throughput can be pushed up to the sensitivity limit of the mass-spectrometry setup, is applicable to any category of protein-ligand interactions and thus bears a wide potential both for high-throughput interactomics and chemoproteomics.

## Introduction

Protein-protein interactions (PPI) are central to cell functioning. Describing cellular interactomes has been the goal of many high-throughput studies over the last few years, resulting in the identification of hundreds of thousands of binary interactions^1^. Different approaches have been used, *eg* yeast 2-hybrid system (Y2H)^2^, affinity-purification mass-spectrometry (AP-MS)^3,4^, protein correlation profiling (PCP)^5^ or proximity-based labeling^6^. Each of these techniques has its own strengths and flaws, resulting in a relatively low overlap between the published interactomes, and a still largely incomplete coverage of the human interactome^1,7–9^. In addition, while PPI affinities can span several orders of magnitude, interactomic data produced by these high-throughput experiments are almost exclusively qualitative (“binds” *vs* “does not bind”). Some approaches were developed that can rank the identified interactions by their affinities, but most are still limited in their throughput and/or in their dynamic range^10–13^. Thus, the accurate description of PPI networks requires new approaches to address interactomes quantitatively, by measuring affinities at a proteome-wide scale.

Many PPI rely on the establishment of interactions between minimal fragments consisting on globular domains and disordered peptide motifs^14–16^. A typical example for such interaction is the 14-3-3 family. The human 14-3-3 proteins consist of seven (β, ε, η, γ, σ, τ and ζ) a-rich homodimeric isoforms encoded by distinct genes that present a high sequence identity^17–19^. 14-3-3s are highly abundant in most human tissues, being systematically found among the top 1% most expressed proteins^20^. They intervene in numerous signaling pathways and cellular processes, ranging from cell cycle progression and apoptosis to signal transduction and protein trafficking. Consequently, they are involved in many pathological conditions, including oncogenic processes, neurodegenerative diseases and viral infections, and are the object of intense drug design research^21,22^. 14-3-3s interact with a plethora of different partners, to such an extent that they constitute one of the largest human PPI networks^3^. This interaction can have many different consequences depending on the targets, including (but not limited to) protein stabilization, hiding or exposure of sub-cellular localization signals or of PPI domains or motifs, chaperone-like activity, and all the changes in intracellular protein trafficking and stability driven by such effects^21^. The 14-3-3 binding site is centered around phosphorylated Ser or Thr residues^23,24^. Additional sequence preferences were described for optimal 14-3-3 binding^23,25^, yet many peptides satisfying this consensus do not bind to 14-3-3 and conversely, many that do not satisfy the same consensus turn out to be *bona fide* interaction partners^26,27^. Finally, the 7 human 14-3-3 paralogs display broadly overlapping sequence binding preferences and protein binding profiles, which raises the question of their specificity^28^.

We previously developed the Holdup assay that quantifies equilibrium binding affinities of PPIs with extremely high precision and dynamic range^9,29–32^, and can reach unmatched experimental throughput when combined with mass spectrometry^33,34^. Here, we demonstrate how the method can be scaled to measure hundreds of protein-peptide affinities at unprecedented accuracy with only one single experiment, by developing and benchmarking a multiplexed version of this assay that uses a label-free quantitative mass-spectrometry strategy. We applied this strategy to the seven human 14-3-3 proteins, and depicted new rules for 14-3-3 binding and specificity, valid at the proteome scale. Our approach is readily transferable to affinity-quantify the interactome of any protein of interest for any pool of ligands.

## Results

### Principle of the Holdup Multiplex

The Holdup Multiplex is a quantitative comparative chromatographic retention assay based on our previously developed Holdup assay^9,29–33,35^, and designed to drastically increase the achievable throughput by quantifying in one single experiment the affinities of thousands of protein-ligand interactions, spanning several orders of magnitude (**Figure 1**). Briefly, a library of *n* different ligands of interest is incubated either with a resin-bound polypeptide (whole protein or fragment), or with an empty, control resin. The total amount of ligand is set up so that the binding is not saturated. After interaction to equilibrium the unbound ligands are filtered-out, without any washing step. The amount of each unbound ligand in the flow-through of the polypeptide sample is then individually resolved in the mix of *n* unbound ligands and quantified relatively to its amount in the flow-through of the control sample, using mass-spectrometry. This relative quantification leads to the dissociation constant of the interaction (*K*_d_), reported as p*K*_d_ values (the negative of the base 10 logarithm of the *K_d_)* for an easier use. Thus in the Holdup Multiplex, the number of interactions quantifiable in one single sample depends on the sensitivity and dynamic reached with the liquid chromatography mass-spectrometry (LC-MS/MS) measurement. Of note is that the ligands with the highest affinities are the most depleted from the unbound flow-through. For that reason, we developed a Data Independent Acquisition (DIA) method to take advantage of a more robust and specific quantification extracted at the MS2 level, and developed ad hoc data analysis procedures (see Methods).

**Figure 1.**
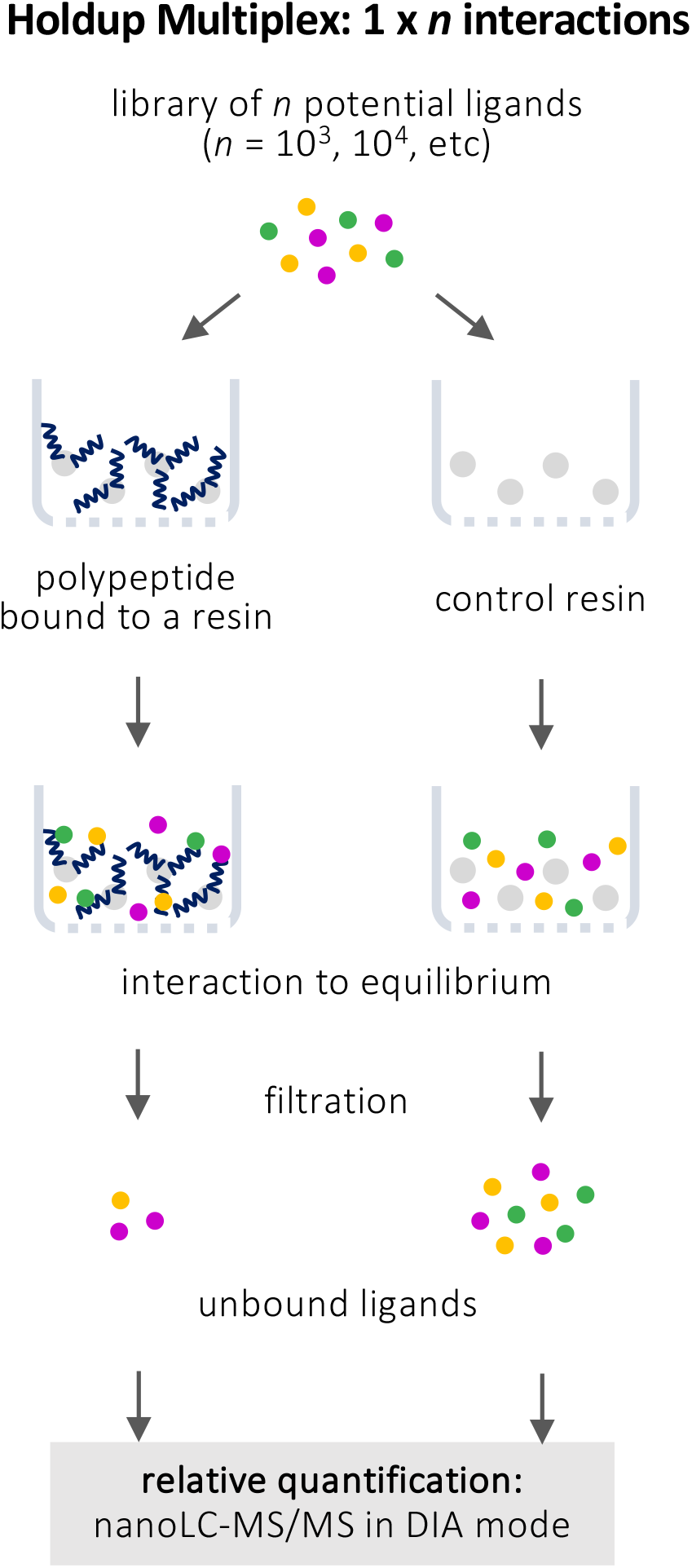
Principe of the Holdup Multiplex. A library of *n* different ligands in solution is applied either to a resin-bound polypeptide, or to an empty, control resin. After interaction to equilibrium and filtration, the amount of each unbound ligand in the flow-through of the polypeptide sample is individually resolved in the mix of *n* unbound ligands and quantified relatively to its amount in the flow-through of the control sample. This relative quantification is performed by nanoLC-MS/MS with a Data Independent Acquisition (DIA) mode, after prior acquisition of a high-quality reference spectral library. The workable size *n* of the library and hence the achievable throughput is limited only by the resolution and sensitivity of the mass spectrometry setup.

### Quantifying the interaction of 14-3-3γ with hundreds of phosphopeptides

We first applied the Holdup Multiplex to study the affinity and specificity of 14-3-3γ (**Figure 2**; see **Table S2** for the detailed results). We designed a library of biologically relevant 14-3-3 binding motifs by crossing the proteins found to interact with several human 14-3-3 in the BioPlex experiment^3^, with the PhosphoSitePlus data to keep only the effectively *in situ* phosphorylated residues^40^. The final library contains 1032 different phosphopeptides, with a third of them following a relaxed definition of the known consensus binding site for 14-3-3, and 90% of them being of unknown 14-3-3 binding status (**Table S1**). Using the Holdup Multiplex we determined the interaction for 14-3-3γ of 659 *ie* 64% of the phosphopeptides of the library (**Figure 2A,B**). Among the remaining phosphopeptides, 86 did not satisfy the different controls set-up for precise quantification and 287 were not detected. An affinity constant (*K*_d_) was measured for 256 *ie* 39% of the 659 determined interactions, with values spanning from 0.7 to 330 μM. The 391 other interactions were weaker than this quantification threshold (*K*_d_ > 330 μM). Finally, a missing-value imputation procedure allowed us to estimate the *K*_d_ of 12 phosphopeptides which 14-3-3 interaction totally depleted from the flow-through. All results are gathered in our ProfAff server (https://profaff.igbmc.science/; see the Method section for more details on the library design, the controls and threshold applied, and the *Kd* estimation procedure).

**Figure 2.**
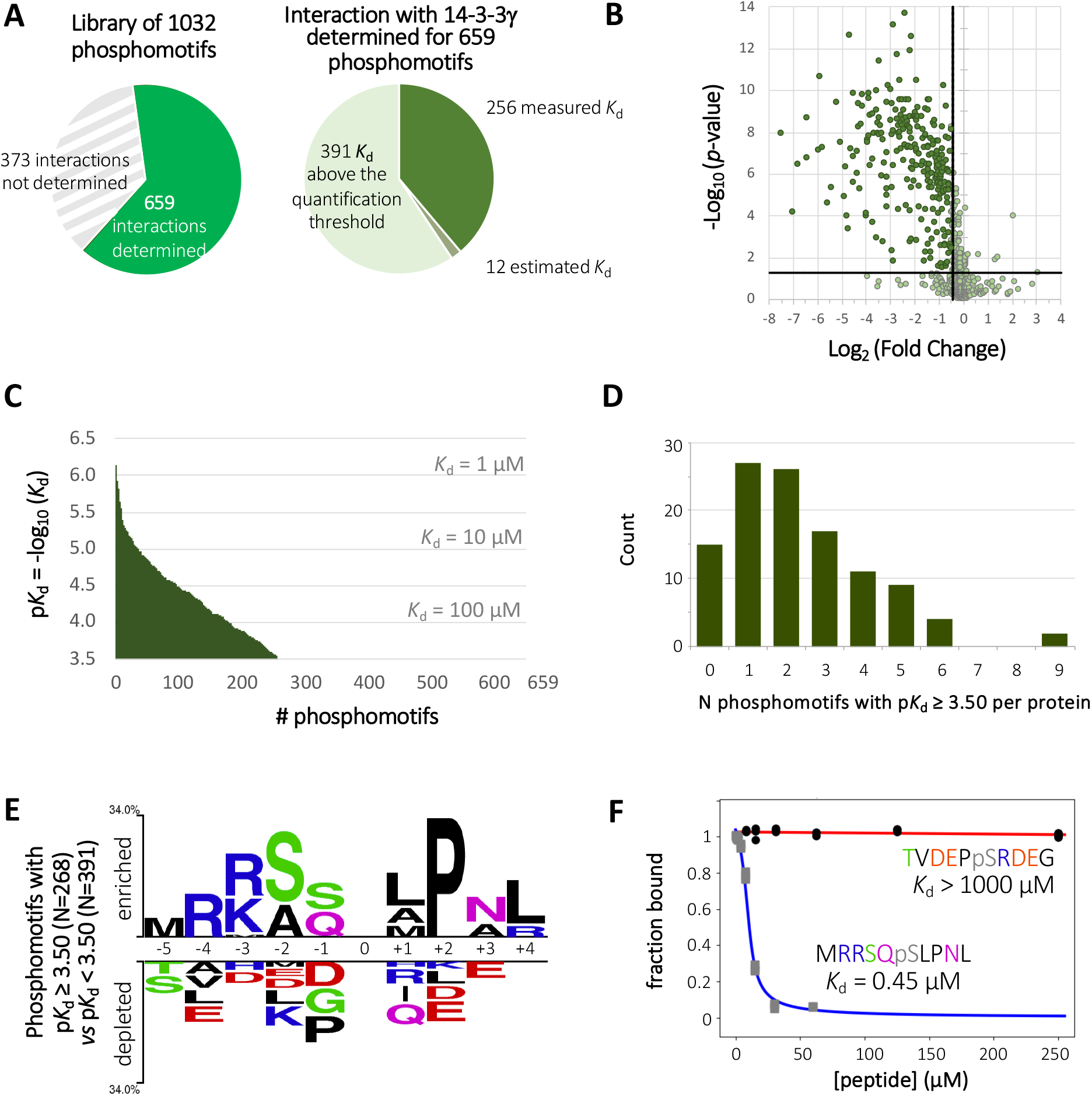
Quantifying the interaction of 14-3-3γ with hundreds of phosphopeptides. **(A)** We determined the interaction with 14-4-4γ for 659 phosphomotifs of the library, and could quantify or estimate a *K*_d_ for 268 of them; the 391 remaining interactions were weaker than the *K*_d_ quantification threshold (*K*_d_ > 330 μM). **(B)** Volcano plot of Holdup Multiplex of the phosphopeptide library against 14-3-3γ. The p-values (p-value ≤ 0.05) and fold-change (Log_2_(FC) ≤ 2*SD{Log_2_(FC)}) thresholds used and are indicated by plain lines. **(C)** Binding profile of 14-3-3γ. The different phosphopeptides are ranked according to their affinity for the protein. **(D)** Number of 14-3-3γ significant binding sites (p*K*_d_ ≥ 3.50) identified per proteins, for the 114 proteins of the BioPlex database selected for this study. **(E)** A sequence logo^43^ for 14-3-3γ binding, calculated over the 659 phosphopeptides which interaction with 14-3-3γ was determined, highlights the sequence differences between phosphopeptides with p*K*_d_ ≥ 3.50 *vs* p*K*_d_ < 3.50. The number of phosphopeptides in each category is indicated within brackets. Position 0 corresponds to the pSer/pThr residue. **(F)** Affinity of 14-3-3γ for phosphopeptides designed according to the positive and negative sequence logos of (E). Affinities were measured by competitive fluorescent polarization. The sequences of the peptides are indicated, with the residues coloured according to the colour-code of the sequence logos. The titrations as well as further examples are available in Figure S2A.

The high proportion of phosphomotifs that significantly interacts with 14-3-3γ (*K*_d_ ≤ 330 μM; p*K*_d_ ≥ 3.50) indicates the high promiscuity, or multi-specificity, of 14-3-3γ. This is further illustrated by its binding profile, in which the different phosphomotifs determined are ranked according to their affinity (**Figure 2C**). In our library, many proteins had different phosphosites assessed. We could identify two or more “functional” binding sites (p*K*_d_ ≥ 3.50) for approximately 60% of the analyzed proteins (**Figure 2D**). Among them, some like CDC25B are well known 14-3-3 binders to which our data add new 14-3-3 binding sites to the previously described ones, all susceptible to participate to the 14-3-3-mediated regulation of the protein^41^. Other like liprin-a1, ALS2 or the NEK1 kinase, were consistently identified as 14-3-3 binders in several orthogonal high-throughput interactomic studies^3,4,42^, yet with no binding site ever identified up to now.

The sequence logo^43^ of 14-3-3γ phosphopeptide binders (p*K*_d_ ≥ 3.50) *vs* non binders (p*K*_d_ < 3.50) highlights sequence preferences inferred from our large scale data (**Figure 2E**). While some sequence logo characteristics correspond to the previously known preferences for 14-3-3 binding (*ie* Arg in −3 and −4), only the over-representation of a Pro residue in +2 of the phosphosite is statistically significant. Clustering using fixed position did not reveal any additional, conditional constraints (not shown). Remarkably, peptides designed according to the positive or the negative logos bound 14-3-3γ with submicromolar affinities or did not show any detectable binding, respectively (**Figures 2F** and **S2A**). Yet, 91% and 59% of the 268 phosphobinders (p*K*_d_ ≥ 3.50) do not follow neither the historical (R-S-X-[pSpT]-X-[PG] for class I; R-X-[YF]-X-[pSpT]-X-[PG] for class II; [pSpT]-X0-2-COOH for class III motifs)^23,25^ nor a more relaxed ([RK]-X_2-3_-[pSpT]-X-[PG] for internal motifs; [pSpT]-X_0-2_-COOH for C-terminal motifs) consensus definition for 14-3-3 binding, respectively. Conversely, as much as 39% of the phosphomotifs following this relaxed consensus do not bind 14-3-3γ significantly (p*K*_d_ < 3.50). Altogether, the large amount of quantitative data obtained highlight how much the sequence preferences for 14-3-3 binding are relaxed, leading to a high multi-specificity.

### The seven 14-3-3 proteins display similar specificities and staggered affinities

Next, we performed and compared the same measurements for the seven human 14-3-3 isoforms (**Figures 3** and **S3**; see **Table S2** for the detailed results). To avoid biases, we used for comparison a subset of 563 phosphopeptides for which binding properties could be determined for the entire 14-3-3ome (**Table S3**). Strikingly, the seven binding profiles were highly comparable, with no reshuffling between the 14-3-3, and no peptide that would be specific of any 14-3-3 (**Figure 3A**). Sequence logos^43^ for the seven 14-3-3 confirmed these comparable behaviors (**Figures 3B** and **S4**). Affinity-weighted frequency logos (in which the sequences of the significant binders are weighted by their affinities^9^) did not allow further discrimination, demonstrating that the same sequence characteristics are required to bind with high affinity to all the human 14-3-3s (**Figure S5**). Thus, the seven human 14-3-3s have very close specificities.

**Figure 3.**
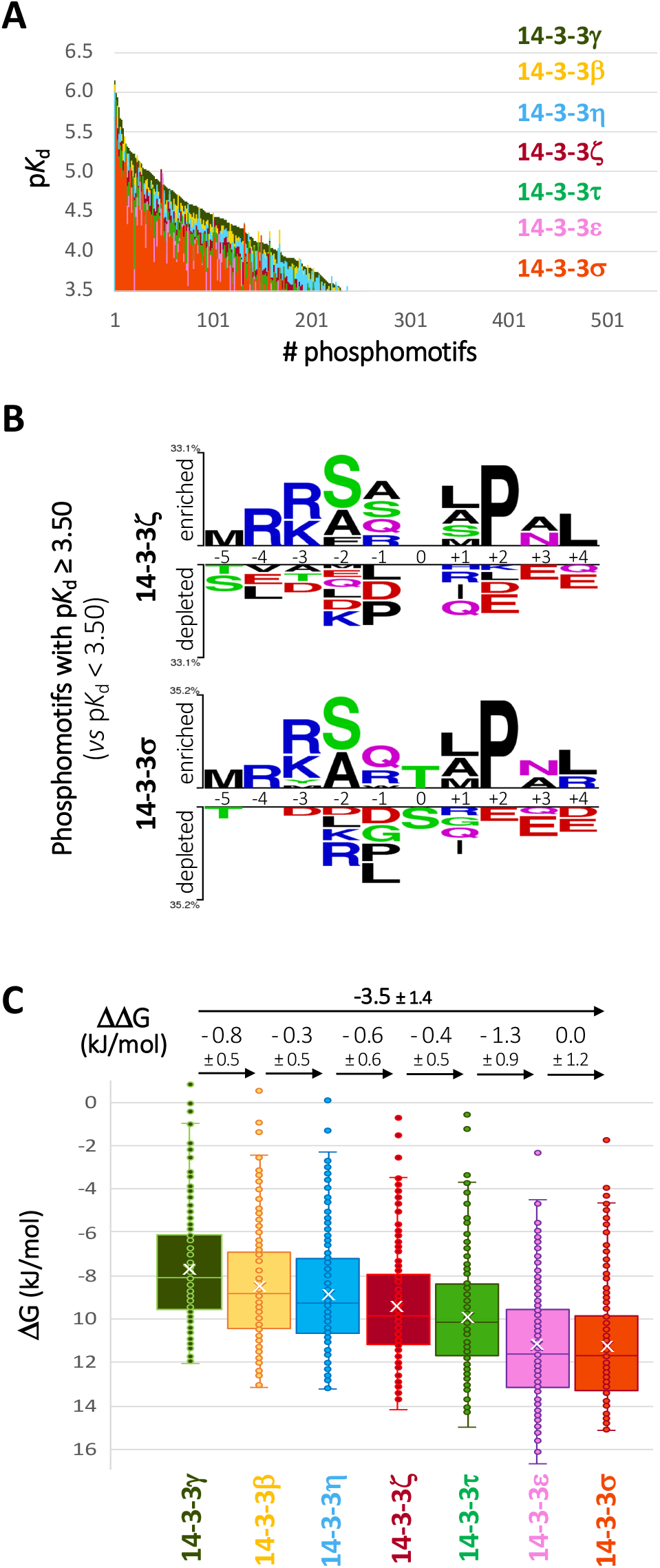
The seven human 14-3-3 isoforms display similar specificities and staggered affinities. **(A)** The binding profiles of the 7 human 14-3-3s for the 563 phosphomotifs which interaction has been determined for all the 14-3-3 are overlaid, with the phosphopeptides ranked according to their affinity for 14-3-3γ. **(B)** Sequence logos^43^ for 14-3-3 ζ and σ binding, calculated over for the 563 phosphomotifs which interaction has been determined for all the 14-3-3. Position 0 corresponds to the pSer/pThr residue. The seven 14-3-3 logos are presented altogether in Figure S4, and the seven affinity-weighted frequency logos^9^ in Figure S5. **(C)** Distributions of the ΔG values of 14-3-3 / phosphomotifs interactions for the different 14-3-3, calculated for the 165 phosphomotifs for which a *K*_d_ was quantified for all the 14-3-3s.

But if the ranking of phosphopeptides presented in **Figure 3A** is similar, their p*K*_d_ are staggered between the different 14-3-3s. A more precise picture is given by the distributions of the binding energies (ΔG), calculated using the 165 phosphopeptides for which a *K_d_* was measured for the entire 14-3-3ome (**Figure 3C**). All 14-3-3s are able to bind each of these 165 phosphopeptides but with different affinities in the following order: γ > β > η > ζ > τ > ε » σ. The average binding energy difference between g (strongest) and σ (weakest) is ΔΔG = −3.5 ± 1.4 kJ/mol, corresponding to a mean *K*_d_ ratio of 5. Surprisingly, only 2 peptides out of 165 did not followed this affinity trend: the phosphopeptide centered on pSer358 from HDAC7 (N°440), which binds to all the 14-3-3 with the similar affinity, and at the opposite the phosphopeptide centered on pSer75 from CDK17 (N°216), which affinities display a marked dependence to the 14-3-3 isoform, with a *K*_d_ ratio of 60 between γ and σ (**Table S3**; see Methods for the outliers identification procedure).

In conclusion, our results demonstrate that the seven human 14-3-3 bind to the same phosphopeptides but with staggered affinities: whatever the phosphopeptide and its affinity for the 14-3-3s, the affinities always follow the same order, with γ having the highest affinity and ε and σ the lowest.

### Assessing phosphopeptide targets of Fusicoccin A at an unprecedented scale

Fusicoccin A (FCA) is a phytotoxin which displays protective activity on mammalian cells in culture through the stabilization of 14-3-3 complexes^44–46^. This stabilization is observed when the side chains of the residues downstream to the phosphoresidue allow the FCA insertion at the interface between the 14-3-3 binding groove and the phosphopeptide docked within the groove^25,28^. This is especially the case with C-terminal phosphopeptides, which display only 0 to 2 residues after the phosphosite^28^. A limited number of interactions between 14-3-3 and internal phosphosites have been reported to be affected by FCA as well (see **Table S4** and references herein).

To determine new 14-3-3 binding sites targeted by FCA, we measured its effect on the interactions between 14-3-3 (γ and σ) and our phosphopeptides library (**Figure 4**; **Tables S2** and **S4**). We identified a total of 135 phosphopeptides which interaction with 14-3-3γ and/or σ was significantly affected by FCA (**Figure 4A**). Most of them were internal to the protein sequences, increasing the amount of known FCA-affected 14-3-3 internal binding sites by a factor of 12. FCA both stabilized and destabilized the interactions between 14-3-3s and their phosphomotifs, in roughly equal proportions (**Figure 4A** and **Table S4**). Remarkably, we observed a very good quantitative correlation between the effect of FCA on 14-3-3γ and 14-3-3σ interactions, with a PCC between the ΔΔG_14-3-3γ_ and ΔΔG_14-3-3σ_ of 0.83 (**Figure 4B**). Finally, we determined that the sequence logos^43^ that favor either the stabilization or the destabilization of the 14-3-3 interaction by FCA, are different (**Figure 4C**). Phosphomotifs with FCA-stabilized interactions present sequence preferences downstream to the phosphorylated residue for small and/or flexible residues, which would favor the accommodation of FCA within the complex^28,47–49^. In contrast, phosphomotifs with FCA-destabilized interactions, which sequence preferences are presented here for the first time, favors the presence of large and/or rigid residues downstream to the phosphosite. In both cases we also observe different sequence constraints upstream to the phosphorylation site, which is unexpected. Peptides designed according to the FCA-stabilized or - destabilized logos present a >2x affinity increase or decrease in the presence of FCA, respectively, further confirming the validity of these consensus (**Figures 4C** and **S2B**). Altogether, these results illustrate how the Holdup Multiplex could be used for compound screening.

**Figure 4.**
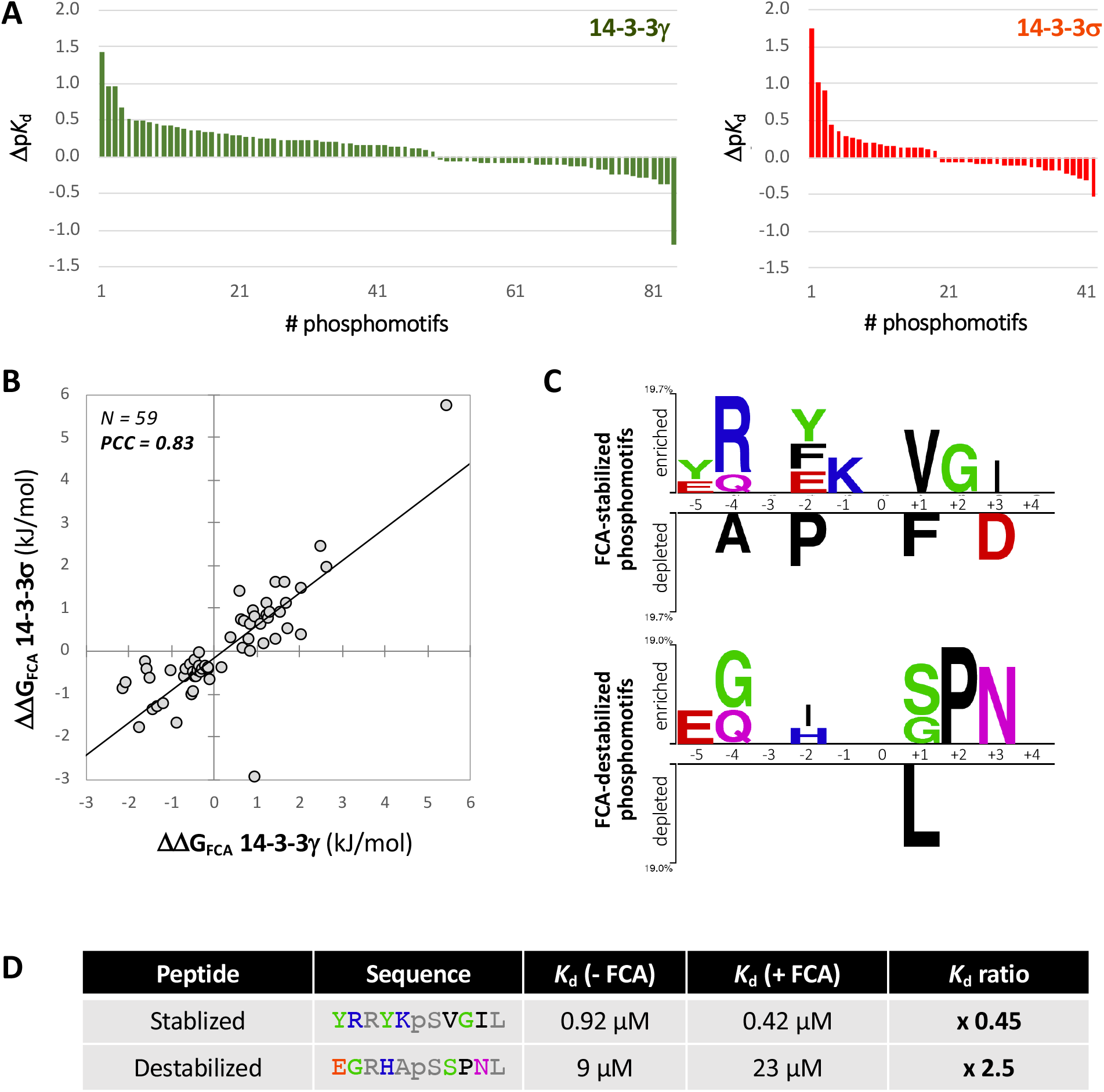
Assessment of 14-3-3 phosphosites targets of Fusicoccin A (FCA) at an unprecedent scale. **(A)** Dp*K*_d_ distributions of the phosphomotifs on which a significant effect of FCA was quantified, for 14-3-3γ (left) and 14-3-3σ (right). A total of 135 significant phosphosites targets of FCA was identified (Table S4), but in some cases Δp*K*_d_ calculation was not possible (one interaction quantified and the other below the quantification threshold). **(B)** Correlation between the effect of FCA on ΔG 14-3-3γ and ΔG 14-3-3σ, for the 59 phosphopeptides on which a significant effect on 14-3-3γ and/or 14-3-3σ was measured. **(C)** Sequence logos^43^ for stabilizing (upper panel) or destabilizing (lower panel) effect of FCA on 14-3-3 / phosphopeptides interactions. The sequences of the 73 and 62 phosphomotifs for which a significant stabilizing or destabilizing effect of FCA on 14-3-3γ and/or 14-3-3σ interaction was measured, respectively, was compared to the sequences of the 145 phosphomotifs for which no effect of FCA neither on 14-3-3γ nor on 14-3-3σ was found. **(D)** Affinity of 14-3-3γ for phosphopeptides designed according to the sequence logos for stabilizing or destabilizing effect of FCA. The sequences of the peptides are coloured according to the colour-code of the sequence logos in (C), and neutral positions are coloured in grey. Affinities were measured by competitive fluorescent polarization. The titrations are available in Figure S2B.

### Performance, evaluation and validation of the Holdup Multiplex

Thanks to different measurements performed throughout this study we extensively tested and validated the Holdup Multiplex (**Figure 5**). Its major advantage is to provide multiple measurements in one single experiment, which both operates a jump in the achievable throughput and facilitates the precise quantitative comparison of thousands of protein-peptide interactions at once. Indeed, we measured, with high precision and in a few sample-runs only, 2,134 unique affinity constants, and determined ≈ 4,000 interactions as weaker than our affinity quantification thresholds.

**Figure 5.**
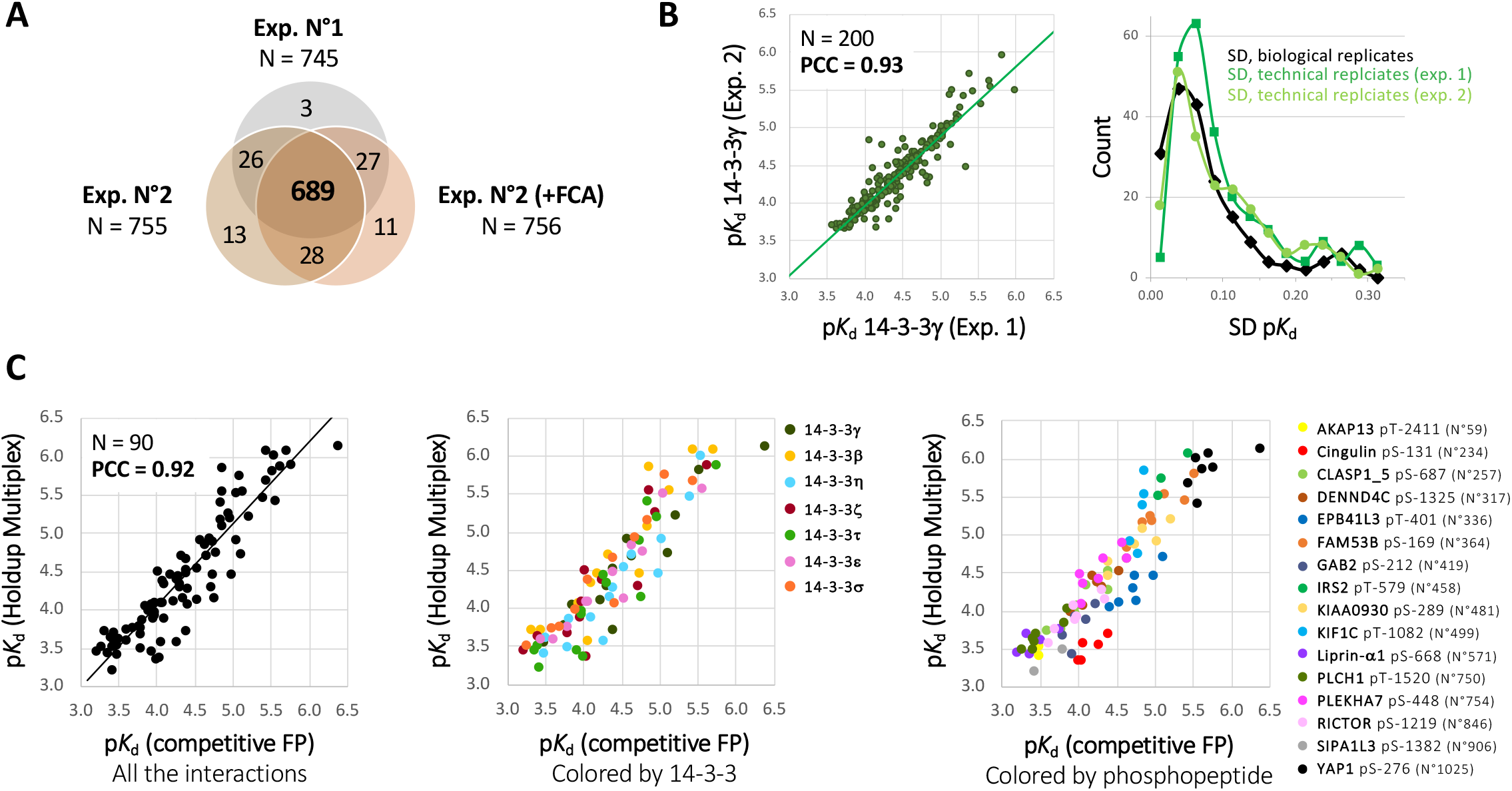
Performance and validation of the Holdup Multiplex. **(A)** Venn diagram of the number of phosphopeptides quantified in the controls of each experiment, showing a high overlap between the 3 independent experiments. **(B)** Reproducibility of the Holdup Multiplex. Left panel, correlation between the p*K*_d_ measured in 2 independent experiments. Right panel, the p*K*_d_ standard deviations between independent experiments (biological replicates) and inferred from one experiment (technical replicates) are similar. Results are shown for 14-3-3γ; results for 14-3-3β, ε and σ are shown Figure S6. **(C)** Orthogonal validation of the Holdup Multiplex. The data are represented with the same colour for all points (left panel), or coloured by 14-3-3 (middle panel) or by phosphopeptides (right panel) to highlight potential biases. For competitive FP, direct and indirect titrations are presented in Figure S7.

We could quantify a large proportion of the 1032 phosphopeptides of the library (73 ± 1 %), with a high overlap of 92% between 3 independent experiments (**Figure 5A**). This very high consistency attests that the 235 phosphopeptides that were not detected (23% of the library) have very likely suffered from poor purity or synthesis yield and/or poor response factors in LC-MS/MS. Indeed, some peptides were particularly long (up to 30 residues) or highly charged, probably leading to too highly charged ion precursors. Others, on the contrary, may have been poorly ionized and/or fragmented because of their non-tryptic nature. However, the main difficulty held by our peptide library was its purity. The synthesis of these phosphorylated, non-tryptic peptides was particularly challenging, and resulted in low and variable purity yields. Considering all these difficulties, the coverage of the tested interactions is very satisfying.

The affinity range achieved by our measurements spans more than 3 orders of magnitude (from 0.7 μM for 14-3-3γ / YAP1 pSer276 interaction to ≈ 900 μM for 14-3-3ε / IRS2 pSer384 interaction – phosphopeptides N°1025 and 452, respectively), which is remarkable. Difficulties may be encountered with high affinity peptides (*K*_d_ < 1 μM), which may be totally depleted from the quantified flowthrough. This can be overcome by adapting the experimental setup (increasing the concentration of protein on the resin and/or of the targeted ligand in the pool). On the other side of the affinity scale, the upper *K*_d_ quantification limit is determined in part by the amount of 14-3-3 bound on the resin (see Methods).

We assessed the precision of the Holdup Multiplex by performing two independent experiments for 14-3-3γ, β, ε and σ. As illustrated in **Figures 5B** and **S6A**, the p*K*_d_ measured by the two experiments were in perfect agreement and highly reproducible (PCC > 0.9 in each case). Moreover, the p*K*_d_ standard deviations between biological and technical replicates were similar (**Figures 5B** and **S6B**).

Finally, we evaluated the accuracy of the affinities measured by Holdup Multiplex. A total of 90 p*K*_d_ between the seven human 14-3-3s and a panel of 16 phosphopeptides of our library, chosen among 16 different protein binders, were measured both by Holdup Multiplex and by competitive fluorescent polarization (FP)^9,28,50^ (**Figures 5C** and **S7**). The quantitative correlation between the p*K*_d_ measured by these two orthogonal approaches over 3 orders of magnitude is remarkable (PCC = 0.92; **Figure 5C**, left panel). The differences between the p*K*_d_ values measured by competitive FP *vs* Holdup Multiplex follow a normal distribution centered on 0 with a standard deviation of 0.3 p*K*_d_. The same quantitative correlation stands for the seven different 14-3-3s considered individually, with no apparent bias (**Figure 5C**, middle panel). However, 2 out of the 16 phosphopeptides tested had all their p*K*_d_ values measured by competitive FP systematically higher than those measured by Holdup Multiplex (phosphopeptides N°234 *ie* pSer131 from Cingulin and N°336 *ie* pThr401 from EPB41L3; **Figure 5C**, right panel). In the Holdup Multiplex measurements, the intensities of those 2 peptides in the control samples were very consistent (coefficients of variation ≈5% in both cases; see **Table S2**). On the other hand, competitive FP affinity measurements rely on a precise estimation of the peptide concentrations used; the later were determined based on the peptide dry weights, and are most probably at the origin of the discrepancy observed between the p*K*_d_ measured by the two methods for these two peptides.

In conclusion, these different results demonstrate the Holdup Multiplex capacity to quantify the interaction between one protein and a library of hundreds of polypeptides of interest simultaneously, with good coverage, a dynamic range of 3 orders of magnitude, and excellent precision and accuracy.

## Discussion

Here, we develop the Holdup Multiplex to simultaneously quantify hundreds of protein-peptide interactions. We assessed the precision and the robustness of this new method and validated it by orthogonal measurements. The range of affinities that can be quantified with accuracy in a single experiment span 3 orders of magnitude, which is unprecedented^34,51–56^. We applied it to quantify the interaction of the seven human 14-3-3 with a library of more than 1000 phosphopeptides, and depicted new rules for 14-3-3 binding and specificity, valid at the proteome scale.

First, we demonstrate the high promiscuity of human 14-3-3, able to bind to a myriad of phosphopeptides over a high range of affinities. We uncovered plethora of new binding sites, more than twice the number of 14-3-3 binding sites identified in the last two decades^26,57,58^. New binding sites were identified both for well-studied 14-3-3 partners or for protein binders only identified through high-throughput interactomic approaches. Each of these interactions will have unique and important biological importance, and our data will be a useful starting point to study them. Most of the characterized 14-3-3 interactions have moderate affinities (10 μM < *K*_d_ < 900 μM), which are hardly detected by classical high-throughput interactomic approaches. Still, they are particularly relevant for 14-3-3 proteins, firstly because such transient interactions are widely used in cell signaling^59^. In addition, 14-3-3 are dimers that can bind cooperatively two different phosphosites on the same protein, and secondary phosphosites of low affinity can synergically increase the affinity of the primary site^60–62^. Finally, the ultimate complex concentration depends not only on the intrinsic affinity between the partners but also on their concentrations. Since 14-3-3 proteins are highly abundant proteins, even the interactions with such moderate affinities can lead to complexes of substantial abundancies.

Our high-throughput peptide-centered approach gives also information on the sequence requirement for 14-3-3 binding, demonstrating their relaxed definition and further underlining the promiscuity of 14-3-3 binding that comes with it. Indeed, a large proportion of the phosphobinders do not follow the relaxed sequence consensus defined by the very same data. Here this proportion is even higher than previously observed^26,27,63,64^, despite a design meant to enrich the library in phosphosites following 14-3-3 binding consensus. Together with their high cellular abundance, the high promiscuity and loose sequence binding requirements of 14-3-3 underline their plethoric and diverse effects on cellular biology.

We performed a large-scale quantitative comparison between the affinities of the seven human 14-3-3 and revealed striking similarities between them. In particular, we could not identify any difference in the specificity of the different human 14-3-3, at least with our peptide library. Previous studies described only restricted numbers of isoform-specific peptide binders^63,65^. Of note, a lack of sequence binding specificity between the human 14-3-3 does not imply a lack of functional specificity: human 14-3-3 have different sub-cellular localizations as well as different expression patterns, both at the tissular level and in response to specific stimuli^20,66,67^. But if 14-3-3s bind to the same peptides their affinities are staggered, with 14-3-3γ being the strongest binder and ε and σ the weakest, confirming on a large scale and with an unbiased library what we previously observed on a discrete set of data^28^. The sequences of the phosphobinding clefts being almost identical between the different 14-3-3, these affinity differences arise most probably from fine conformational effects spanning the entire structure^18,28^. Altogether, these properties suggest that the seven 14-3-3 isoforms might collectively serve as a redundant buffer of highly phosphorylated proteins during intracellular phosphorylation boosts, as previously hypothesized^28,68^.

Fusicoccin-A (FCA) is a widely studied phytotoxin that display anti-apoptotic and neuroprotective effects in mammalian cells. This small molecule has been described to stabilize the interactions of 14-3-3 with some of its partners, and is a model to study PPI modulation^69^. We quantified the effect of the phytotoxin Fusicoccin-A (FCA) at an unprecedented scale, uncovering dozens of potential new FCA targets. Contrarily to what has been observed previously on a limited set of peptides^70^, we could demonstrate the absence of isoform-specificity effect of FCA, at least between 14-3-3γ and σ. Unexpectedly, many phosphosites which interaction was affected by FCA were internal to the protein sequences, and both stabilization and destabilization effects of FCA on the interactions were observed. Among the newly identified FCA targets, some of them like YAP1 or RICTOR are at the center of important signaling pathways; the stabilization or destabilization of their interaction with 14-3-3 proteins may have pleiotropic effects. Thus, the effect of FCA on the interaction between 14-3-3 and their cellular targets may be more massive than previously thought, explaining the complexity of the physiological effects of this molecule^45,46,71^.

Finally, our results further demonstrate the versatility of the Holdup assay, applied here to yet another protein system. Actually, the Holdup Multiplex could be applied to any kind of PPI involving motifs, domains or full-length proteins, provided that the polypeptides of the library can be individually quantified by mass-spectrometry. Alternatively, full-length protein binders can be quantified by massspectrometry from cellular extracts^33^. Here, we used a library of 1000 peptides, but in practice the achievable throughput is only limited by the LC-MS/MS setup, which is routinely able to quantify tens of thousands of peptides in classical bottom-up proteomics experiments. Furthermore, we anticipate that mass spectrometry-based quantification of non-phosphorylated synthetic peptides or of trypsinable domains or proteins will lead to higher recovery rates than the ones obtained here with non-trypsic phosphopeptides. Pushing the principle further, the Holdup Multiplex could be applied to quantify the interaction of any kind of resin-attachable molecule with a library of any kind of quantifiable ligands, and thus to the interactome quantification of nucleic acids, sugars, lipids or small molecules, with countless applications not only for system biology but also for drug research and screening.

## Material and Methods

### Human 14-3-3 cloning and purification

The different constructs used in this study and their sequence information are described in **Figure S1.** All the constructs were validated by DNA sequencing, and the identity and sequences of the purified proteins were assessed by mass spectrometry peptide mapping. For the Holdup experiments, AviTag-His_6_-MBP-TEVsite (used as negative control) and the seven AviTag-His6-MBP-TEVsite-14-3-3 constructs were cloned into pET bacterial expression vectors. The proteins were expressed in E. coli BL21(DE3) with IPTG induction (1 mM IPTG at 25°C for 4 hours) and harvested cells were lysed in a buffer containing 50 mM Tris-HCl pH7.5, 150 mM NaCl, 2 mM β-mercapto-ethanol, cOmplete EDTA-free protease inhibitor cocktail (Roche, Basel, Switzerland), and trace amount of DNAse, RNAse, and lysozyme. Lysates were frozen at −20°C before further purification steps. After thawing lysates were sonicated and centrifuged for 1h for clarification. Expressed proteins were captured on pre-packed Ni-IDA (Protino Ni-IDA Resin, Macherey-Nagel, Duren, Germany) columns, and washed with 10 column volumes of cold wash buffer (50 mM Tris-HCl pH7.5, 150 mM NaCl, 2 mM β-mercapto-ethanol) before elution with 250 mM imidazole. The Ni-elution was collected directly on a pre-equilibrated amylose column (amylose high flow resin, New England Biolabs, Ipswich, Massachusetts). Amylose column was washed with 5 column volumes of cold wash buffer then eluted in a buffer containing 25 mM Hepes pH 7.5, 150 mM NaCl, 1 mM TCEP, 10% glycerol, 5 mM maltose, cOmplete EDTA-free protease inhibitor cocktail. The concentration of proteins was determined by their UV absorption at 280 nm before aliquots were snap-frozen in liquid nitrogen and stored at −80°C. MBP-fused 14-3-3 isoforms used for fluorescence polarization and their purification procedure are described in ^9^.

### LC-MS/MS peptide mapping to validate the 14-3-3 protein sequences

The AssayMAP Bravo platform (Agilent Technologies) was used to perform protein cleanup and in-solution digestion of the seven 14-3-3 isoforms. Briefly, proteins were desalted using solid phase extraction C18 cartridges (Agilent, 5 μL bed volume). Then, the proteins were reduced with 10 mM DTT for 1 h at 37 °C and then alkylated with 20 mM iodoacetamide for 30 min in the dark at RT. Next, the samples were digested with Trypsin in a 1:25 w/w ratio (enzyme/protein) in 50mM ammonium bicarbonate and incubated at RT for 4h. After cup wash and internal cartridge wash with 0.1% TFA, trypsic peptides were eluted with 0.1% TFA in 70% ACN at 5μL/min. Eluted peptides were dried and resuspended in 0.1%FA in H2O.

NanoLC-MS/MS analyses were performed on a nanoACQUITY Ultra-Performance LC system (Waters, Milford, MA) coupled to a Q-Exactive Plus Orbitrap mass spectrometer (ThermoFisher Scientific) equipped with a nanoelectrospray ion source. The solvent system consisted of 0.1% formic acid in water (solvent A) and 0.1% formic acid in acetonitrile (solvent B). Samples were loaded into a Symmetry C18 precolumn (0.18 x 20 mm, 5 μm particle size; Waters) over 3 min in 1% solvent B at a flow rate of 5 μL/min followed by reverse-phase separation (ACQUITY UPLC BEH130 C18, 200 mm x 75 μm id, 1.7 μm particle size; Waters) using a linear gradient ranging from 1% to 35% of solvent B at a flow rate of 450 nL/min. The mass spectrometer was operated in DDA mode by automatically switching between full MS and consecutive MS/MS acquisitions. Survey full scan MS spectra (mass range 300-1800) were acquired in the Orbitrap at a resolution of 70K at 200 m/z with an automatic gain control (AGC) fixed at 3.10^6^ and a maximal injection time set to 50 ms. The ten most intense peptide ions in each survey scan with a charge state ≥ 2 were selected for fragmentation. MS/MS spectra were acquired at a resolution of 17,5K at 200 m/z, with a fixed first mass at 100 m/z, AGC was set to 1.10^5^, and the maximal injection time was set to 100 ms. Peptides were fragmented by higher-energy collisional dissociation with a normalized collision energy set to 27. Peaks selected for fragmentation were automatically included in a dynamic exclusion list for 60 s. To minimize carry-over, three solvent blank injections were performed after each sample.

The nanoLC-MS/MS.raw files were converted to peaklist.mgf files and searched with the Mascot search algorithm (local Mascot server, version 2.5.1, Matrix Science, London, UK) against a custom protein database containing the sequences of the seven 14-3-3 proteins. Full trypsin specificity was set. Parent and fragment mass tolerances of 10ppm and 0.07Da, respectively were defined. Carbamidomethyl (C) was set as a fixed modification, Oxidation (M) was set as variable modification.

### Peptides and peptide library

#### Design of a library of 1032 potential 14-3-3 phosphomotifs

To design the most biologically relevant 14-3-3 phosphomotif library, we implemented a strategy that combines both the BioPlex ^3^ and PhosphoSitePlus ^40^ databases. We started from the common 114 prey proteins captured by 5 different 14-3-3 isoforms in the AP-MS BioPlex experiment (γ, β, η, τ and ζ; ε was not tested as bait and σ fished almost no proteins apart from the other 14-3-3 isoforms). Then we used PhosphoSitePlus (version January, 2021) to assess the presence of *in situ* phosphorylated Ser or Thr residues in these 114 proteins. All the phosphosites present in PhosphoSitePlus and corresponding to a loose 14-3-3 binding consensus definition ([R/K]-X_2-3_-[pS/pT]-[^P]-[P/G] for internal motifs, and [pS/pT]-X_0-2_-COOH for C-terminal motifs) were kept (301 phosphosites). This definition does not correspond to the historical 14-3-3 binding motif definition (R-S-X-[pS/pT]-[^P]-[P/G] for class I and R-X-[Y/F]-X-[pS/pT]-[^P]-[P/G] for class II internal motifs) ^23^. Indeed, it was chosen to be as much inclusive as possible, considering that many experimentally characterized 14-3-3 motifs do not follow these classical motif definitions^26,27^. Only 70 of these 301 phosphosites obeyed to the classical 14-3-3 binding motif.

For the remaining phosphosites not following this loose 14-3-3 binding consensus definition, the following filtration steps were applied:

All the phosphosites identified by at least 1 low throughput paper in the PhosphoSite Plus database (LTP; determined using methods other than discovery mass spectrometry) were kept.
All the phosphosites identified by at least 10 high throughput papers in the PhosphoSite Plus database (HTP; determined using only proteomic discovery mass spectrometry) were kept. This threshold was set up because high-throughpout phosphoproteomics data are not always reliable when considered individually^72^. Of note, a recent study observed that 14-3-3 binding sites are often more frequently phosphorylated in phosphoproteomic studies^57^.
If after these steps the number of selected phosphosites for one protein was <3, we included phosphosites that do not follow the above-described rules, until reaching at least 3 phosphosites / protein.

10-mer phosphopeptides were obtained from the selected phosphosites by extending the natural protein sequence in N-terminal (5 residues) and C-terminal (4 residues) directions, with some exceptions. When the resulting 10-mer peptide had a Cys at the C-terminus this design was shifted of 1 residue to avoid potential difficulties at the synthesis step. For C-terminal phosphosites the N-terminal sequence was extended until the total peptide length reached 10 residues. After suppression of duplicates the final number of phosphomotifs designed with this approach was 1008.

Finally we completed this library with 24 peptides for which affinity measurements were previously collected for all the human 14-3-3ome (see **Table S1.2** and references herein). The final library of 1032 phosphopeptides is described in **Table S1**, and contains 927 phosphosites which binding to 14-3-3 was never characterized.

#### Peptides synthesis and preparation

Most of the peptides composing the peptide library was synthetized as crude peptides by JPT Innovative Peptide Solutions (https://www.jpt.com/) with a Spot-technology. Mass-spectrometry analysis on 53 peptides (5% of the library) confirmed their presence and assessed their purity, which was between 6 and 50%, with a mean purity of 20%. Peptides were dissolved in ultra-pure H2O + 1 mM TCEP. Average peptide concentrations were determined based on the mean dry weight and purity provided by JPT. Peptides of the library containing two phosphorylation sites (from CFTR, TASK1 and BLNK – phosphopeptides N°1, 2 and 3, respectively) as well as the peptides used for fluorescence polarization were chemically synthesized on an ABI 443A synthesizer with standard Fmoc strategy and HPLC-purified by the peptide synthesis service at IGBMC (https://www.igbmc.fr/en/plateforms-and-services/platforms) with >95% purity. Peptide concentrations were determined based on their dry weight.

### Fundamentals of the Holdup assay

#### General principle

The Holdup assay is quantitative interactomics assay developed and extensively validated over the last decade by our team^9,29–32,35–37^. It can measure protein-peptide affinities at unprecedented throughput and accuracy, and was successfully applied to measure the affinities of tens of thousands of domain-motif interactions, including PDZ-PBM complexes^9,29,39^, E6-LxxLL motif complexes^31^ and RCC1-like domain-peptide complexes (unpublished results). All these results are available on our dedicated served ProfAff (https://profaff.igbmc.science/), an on-line tool to store, display and analyze our quantitative interactomic data at proteome-wide scale.

The Holdup assay is a comparative retention-based chromatographic approach devoid of washing steps, that attempts to monitor the steady-state binding equilibrium by separating the interaction partners on different, moving *vs* stationary, phases (**Figure 1A**, left panel). A resin is saturated by either a control or the polypeptide under study. Then, both resins are incubated with a ligand-containing solution until equilibrium. The unbound ligand in the control and the polypeptide samples are filtered-out and quantified by appropriate methods (see below). Binding intensities (*BI*) are calculated using Eq. 1:

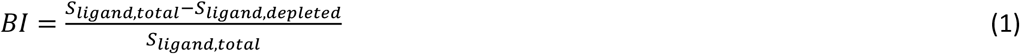

where *S_lignad,total_* corresponds to the total signal of ligand (quantified in the control flow-through) and *S_ligand,depleted_* the signal of unbound ligand (quantified in the flow-through of the polypeptide-saturated resin).

The law of mass action allows *BI* to be converted into a dissociation constant using Eq. 2:

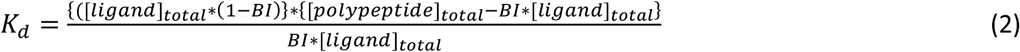

[polypeptide]_total_, the total concentration of polypeptide on the resin, is accessible through different approaches (see below). In some instances [ligand]_total_ (the total ligand concentration applied to the resin) is known and *K*_d_ can be determined using Eq. 2. In other cases [ligand] is not known with precision but can be chosen so that [polypeptide]_total_ >> [ligand]_total_, and Eq. 2 can be approximated with Eq. 3:

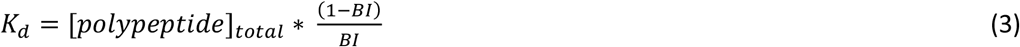

Finally, one can find more convenient to work with log-transformed *K*_d_ values, *e.g*. p*K*_d_ or ΔG, obtained through Eqs. 4 and 5:

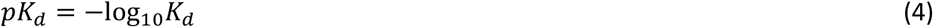

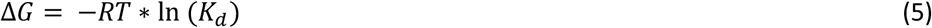

#### General considerations on the affinity range accessible by the Holdup assay

The range of the quantifiable *BI* directly depends on the methods used to quantify the ligand in the flow-through and its performance: highly sensitive methods will be able to quantify very low *S_ligand,depleted_* values, corresponding to high-affinity interactors; precise methods will be able to discriminate between slightly different *S_ligand,total_* and *S_ligand,depleted_* values, corresponding to low-affinity interactors. In addition, the dissociation constant depending directly on the concentration of polypeptide on the resin ([polypeptide]_total_; Eq. 3), the highest amount of polypeptide one can saturates the resin with, the highest *K*_d_ values the Holdup assay will be able to quantify. But a high [polypeptide]_total_ is a double-edged sword, since it will also deplete the highest-affinity binders to an extent at which *S_ligand,depleted_* may drop below the sensitivity threshold of the detection method, preventing its quantification. In summary, both the method chosen for quantification and the set-up of the Holdup experiment ([polypeptide]total) will have an influence on the *K*_d_ range quantifiable by the assay. The different step-up used up to now, described in the following paragraph, measured affinities with high accuracy between 1 and 300 μM.

### Previously developed versions of the Holdup assay and their achievements

Up to now the Holdup assay was performed with a biotinylated peptide motif bound to a streptavidin resin. A resin saturated with biotin was used as a control^29^. Direct determination of the peptide concentration on the resin ([polypeptide]_total_ in Eq. 2 and 3) was not possible. For some selected peptides we measured a set of *K*_d_ by an orthogonal method, then used Eq. 3 to trace back [polypeptide]_total_ from the measured *BI*. A generic [polypeptide]_total_ was obtained by averaging the [polypeptide]total obtained with different peptides^9,30^. Streptavidin resins have a relatively limited binding capacity, and the average [polypeptide]total obtained after resin saturation was 18 μM^9^.

We successfully used different detection methods for measuring the concentration of ligand in the flow-throughs. It is possible to apply to the polypeptide-saturated resin total bacterial lysates over-expressing a domain of protein of interest, which is subsequently quantify in the flow-throughs by capillary electrophoresis^29^. While the ease of use of such a complex matrix is desirable (ease of library preparation, crowded environment, etc.), the readout requires a tedious multi-step data-curation^32^. Alternatively one can used as ligand purified polypeptides and quantify them through their intrinsic fluorescence^31^. Depending on the quality of the purification, this method can reach a very high *BI* sensitivity^9^.

Through the years our Holdup measurements were extensively benchmarked using different orthogonal affinity measurements. Qualitative comparisons with interactions probed by yeast 2-hybrid system, luciferase complementation assay or SPOT peptide array first validated the method^29,31^. We also performed more than 600 *K*_d_ measurements either by split-Nanoluciferase Protein Complementation Assay, by SPR or by competitive fluorescent polarization, and the remarkable quantitative correlations between the dissociation constants measured by Holdup and by these orthogonal techniques demonstrated the accuracy of the Holdup assay^9,29,31,38^. In addition, we observed quantitative correlation between the Holdup measurements obtained on domain-motif interactions, and the enrichment of full-length proteins from cellular extracts in AP-MS experiments^9,33^.

### The Holdup Multiplex

#### Principle of the Holdup Multiplex

The current Holdup assay as presented in **Figure 1A** is parallelizable on multiwell plates and we recently pushed its throughput at its maximum^9^. Thousands of interactions can be measured in a few days by an experienced team. However, the quantitative characterization of interactomes at a proteome level would require to further increase this throughput by several orders of magnitude. With the objective of developing a quantitative interactomics method that could reach this throughput, we implemented the Holdup Multiplex. The basic principle of the Holdup is the same, but instead of having 1 ligand in solution we have a mix of *n* ligands, whose relative amounts in the flow-throughs are individually determined by label-free nanoLC-MS/MS (**Figure 1B**).

#### Seven 14-3-3 human proteins x 1032 potential phosphomotifs

In this article we set up this Holdup Multiplex assay by characterizing the interaction of the seven human 14-3-3 proteins with a pool of 1032 potential 14-3-3 phosphopeptide binders. 150 μL Ni-sepharose resin (Ni Sepharose 6 Fast Flow, Cytiva) was incubated with 200 nmoles of either purified His_6_-MBP (control) or purified His_6_-MBP-14-3-3 in Holdup buffer (50 mM Tris-HCl pH8.0, 300 mM NaCl, 1 mM TCEP) for 2h at 4°C upon gentle mixing, then extensively washed before resuspension in 2 mL Holdup buffer, resulting in a 13x dilution of the resin. We demonstrated in a previous work that the MBP fusion had no influence on the 14-3-3 affinities measured^28^. The His_6_-MBP or His_6_-MBP-14-3-3 concentrations on the resin were determined by pipetting 3×200 μL of diluted resin (corresponding to 3×15 μL of resin) into a filter plate (Millipore, Burlington, Massachusetts), filtering out the buffer by centrifugation (500g), eluting the proteins with 500 mM imidazole before measuring their UV absorption at 280 nm. Then, the His_6_-MBP and His_6_-MBP-14-3-3 resins were diluted in unbound resin so that the different protein concentrations were similar (between 100 and 150 μM in a 45 μL reaction volume, depending on the experiment).

In a second step, the different resins 13x-diluted and loaded with similar protein concentrations were pipetted (5×200 μL, corresponding to 3×15 μL of resin and resulting in 5 technical replicates for each condition) into a filter plate, filtered out and washed by 200 μL Holdup buffer. Each well was then incubated with 30 μL of a mix containing the 1032 peptides of the library diluted in Holdup buffer. The final total peptide concentration in the reaction volume (15 μL resin + 30 μL peptide solution) was set at 10 μM (corresponding to a mean concentration of each peptide of the mix of ≈ 10 nM), so that the potential simultaneous binding of all the peptides would not be saturating. Incubation was carried out for 15 min with shaking. A final filtration step (5 min at 1500g) collected the flow-throughs in nonbinding PCR plates. The 30 μL filtrates were kept at 4°C until treatment for mass-spectrometry.

In a final step the amount of His_6_-MBP or His_6_-MBP-14-3-3 effectively present in each well was determined by eluting the proteins with 500 mM imidazole and measuring their UV absorption at 280 nm. The mean corresponding concentrations in the 45 μL reaction volume calculated over the different replicates and their associated standard error values are listed in **Table S2** and were used for *K*_d_ calculation (Eq. 3).

#### Effect of Fusicoccin-A on 14-3-3 /phosphopeptides interactions assessed by Multiplex Holdup

A 100 μM FCA solution was prepared by first dissolving FCA (gift of NN. Sluchanko) at 5 mM in ethanol, then speed-vac and resuspend it in Holdup buffer. A Holdup assay was performed as described above, with the following modifications. After being displayed in the filter plate the His_6_-MBP or His_6_-MBP-14-3-3 loaded resins were incubated with 50 μL of FCA at 100 μM for 20 min with shaking, then filtered out. The peptide mix incubated was prepared at a total concentration of 10 μM in 100 μM FCA before incubation with the protein-loaded, FCA-equilibrated resins.

### Quantitative analysis of Multiplex Holdup by nanoLC-MS/MS

#### Peptide cleanup

Eluted peptides were desalted using C18 cartridges (5 μL bed volume) on a Bravo AssayMAP Platform (Agilent Technologies). Briefly, C18 cartridges were primed with 100 μl of 70% of acetonitrile with 0.1% TFA, equilibrated with 50μL of 0.1%TFA and loaded with 30 μl of digests. After cup wash and internal cartridge wash with 0.1% TFA, peptides were eluted with 30μl of 0.1% TFA in 70% ACN at 5μL/min. The samples were dried and resuspended in 9 μl H2O, 2% ACN, 0.1%FA and 1 μl of the indexed Retention Time solution (iRT from Biognosys).

#### nanoLC-MS/MS analysis

Data-dependent acquisition (DDA) and data-independent acquisition (DIA) analyses were performed on a NanoAcquity UPLC device (Waters) coupled to a Q-Exactive HF-X mass spectrometer (Thermo Fisher Scientific, Bremen, Germany). Solvent system consisted of 0.1% FA in H2O (solvent A) and 0.1% FA in ACN (solvent B) (Optima LC/MS grade solvents, Fisher Chemical, Illkirch, France). Peptides were loaded onto a Symmetry C18 precolumn (20 mm × 180 μm, 5 μm diameter particles; Waters) over 3 min at 5 μL/min with 1% solvent B. Peptides were eluted on a Acquity UPLC BEH130 C18 column (250 mm × 75 μm,1.70 μm particles; Waters) at 400 nL/min with the following gradient of solvent B: linear from 1% to 8% in 2 min, linear from 8% to 35% in 77 min, up to 90% in 1 min, isocratic at 90% for 5 min, down to 2% in 1 min, and isocratic at 2% for 16 min.

For DDA analyses, full-scan MS spectra were collected from 300-730 m/z at a resolution of 120,000 at 200 m/z with an automatic gain control target fixed at 3.10^6^ and a maximum injection time of 60 ms. The top 20 precursor ions with an intensity exceeding 1e^5^ and charge states ≥ 2 were automatically selected from each MS spectrum for fragmentation by higher-energy collisional dissociation (normalized collision energy set to 27), excluding unassigned, monocharged and over seven times charged ions. Spectra were collected from 200-2,000 m/z at a resolution of 15,000 at 200 m/z, an automatic gain control target fixed at 1.10^5^ and a maximum injection time of 60 ms. Dynamic exclusion time was set to 10 s.

For DIA analyses, full-scan MS spectra were collected from 300-890 m/z at a resolution of 60,000 at 200 m/z with an automatic gain control target fixed at 3.10^6^ and a maximum injection time of 60 ms. Fragment analysis (MS2) was subdivided into 43 windows of 10 m/z widths at a resolution of 15,000 at 200 m/z, an automatic gain control target fixed at 1.10^6^ and an automatic maximum injection time.

#### Reference Spectral Library Generation

Forthy-one DDA and 5 DIA analyses ran on 10 subpools of 96 and 1 subpool of 72 synthetic peptides were used to build a reference spectral library in Spectronaut (v.15.0; Biognosys). Searches were ran against a FASTA database containing the 1032 phosphopeptides’ sequences and the iRT peptides sequences. Oxidation of methionines was set as a variable modification. NoCleavage was used as digestion enzyme. Data was extracted using dynamic mass tolerances. Identification was performed using 10% precursor q-value cutoff.

#### DIA data interpretation

DIA data interpretation was performed in Spectronaut (v.15.0; Biognosys) using the following settings and the same FASTA file as described above. Data was extracted using dynamic mass tolerances. Identification was performed using 10% precursor q-value cutoff. Quantification was performed using interference correction and at least three fragment ions were used per peptide. Quantity values extracted correspond to MS2 XIC peak areas.

#### Data availability

Raw LC-MS/MS data files, the FASTA database as well as the Spectronaut method including full list of parameters have been deposited to the ProteomeXchange Consortium via the PRIDE^73^ partner repository with the dataset identifier PXD035116.

### Mass spectrometry data analysis and inference of dissociation constants

#### Data filtration

Each phosphopeptide detected was possibly present in the form of different precursors: with different charge states (mainly 2+ and 3+) and/or oxidized or not on different residues. We first eliminated all oxidized ions. Indeed, we had no mean to decipher whether the oxidation occurred before or after the Holdup interaction assay, and oxidation of residues flanking the phosphosites could affect the interaction with 14-3-3. Actually, for some peptides we observed a marked reduction of the binding intensity (*BI*) of the oxidized *vs* non oxidized precursors. Of note, this filtering step eliminated very few information; for example, in experiment N°1, 7 phosphopeptides were present only in the form of oxidized precursors. Then, we kept only the precursors that were quantified at least twice in the control, and for which the mean intensity in the control was above 10.000. Finally, when different charge states were present after these two filtering steps, we kept only the most abundant precursor. In the overwhelming majority of cases, the *BI* of precursors with different charge states were similar; when the *BI* differed, the measurement was always more precise (decreased SD and p-values) with the most intense precursor (data not shown).

After these different filtering steps, the total number of phosphopeptides quantified in the controls was 745 (72% of the library) for the 1^st^ experiment, and 755 or 756 (73% of the library) for the 2^nd^ experiment in the absence or in the presence of FCA, respectively. In each case, 91-92% of these phosphopeptides were also quantified in the controls of the other experiments (see **Figure 1B** and main text for more details).

#### Data normalization

Many phosphopeptides of the library effectively bound the resin-immobilized 14-3-3, so a global, sample-based normalization was not possible. Therefore, we developed a normalization procedure adapted to our dataset. We identified an ensemble of non-binding phosphopeptides by the following strategy, and used them as internal standards. We reasoned that by plotting the mean intensities of the precursors in a control *vs* in a 14-3-3 sample, non-binding phosphopeptides would align on a straight-line which slope would depend on the normalization factor to be applied. Thus, we selected precursors with low binding intensities (*BI* before normalization < 5^th^ percentile for 14-3-3γ, β, η and ; > −0.5 and < 6^th^ percentile for 14-3-3τ, ε and σ) and reliable intensity measurements (mean intensity in the control > 3^rd^ percentile; variation coefficients of the intensities in the control and the 14-3-3 sample both < 5^th^ percentile). At least 86 phosphopeptides were selected as internal standards depending on the samples, with a mean of 103 phosphopeptides. For each sample *s* (control or 14-3-3) and each internal standard *i* we calculated a normalization factor N_*s,i*_ so that the intensity of the internal standard in the sample *s* is equal to its mean intensities in all samples (all control and 14-3-3 replicates). Then, for each sample *s* we applied to each phosphopeptide (including the internal standards) a global normalization N_*s*_ that averages all the N_*s,i*_. This normalization step was performed for each 14-3-3 individually. Thus, the normalized intensities of the precursors in the control replicates reported in **Table S2** may vary from one 14-3-3 to another within the same experiment, even if the same control was used.

#### Thresholds for FC and t-tests

Both a *p*-value threshold (calculated by two-tailed unpaired T-test; p<0.05) and a fold-enrichment threshold (corresponding to 2x the standard deviation of the Log_2_(FC) distribution) were applied to identify the phosphopeptides binding to the 14-3-3 with a quantifiable dissociation constant. Again, in each sample too many phosphopeptides bound to the 14-3-3 to consider the Log_2_(FC) distribution as a whole for its SD determination. We reasoned that the positive Log_2_(FC) values (right arm of the volcano plot) were representative of Log_2_(FC) values for non-binders phosphopeptides, whereas negative Log_2_(FC) values (left arm of the volcano plot) were “contaminated” by significant values corresponding to phosphopeptide binders. Thus, the SD of Log_2_(FC) distribution was calculated considering only positive Log_2_(FC) values and their negative mirror values, for each 14-3-3 in each independent experiment. The sensitivity of the interaction detection, *ie* the maximal affinity constant measured, varied between 220 and 900 μM (see **Table S2**). The volcano plots are presented in **Figures 2B** and **S3**.

The remaining phosphopeptides were categorized as follow. A *BI* < −0.5 (FC > 2) was considered as an artefact and the corresponding interactions were classified as “non determined” (n.d.). Interactions for which the FC fulfilled the threshold but not the p-value were also considered as “non determined”. Finally, interactions with *BI* comprised between −0.5 and the binding threshold were considered to be weaker than the quantification threshold; they are annotated as “non-binding” (N.B.) in **Table S2**, **S3** and **S4** for the sake of simplicity.

#### Conversion of Holdup binding intensities to dissociation constants and error propagation

As detailed above, for each 14-3-3 / phosphopeptide interaction a dissociation constant *K*_d_ can be calculated based on the binding intensity and the 14-3-3 concentration (Eq. 3). For each interaction, the standard deviation was propagated from the variability of the intensities measured in the different control and 14-3-3 replicates, to the *BI* and then *K*_d_, the latter by combining it with the SD of the 14-3-3 concentration measurements. Mathematically, the *K*_d_ variability can be further propagated to the p*K*_d_, the ΔG or the ΔΔG values, but the logarithmic relationship between the former and the latter renders these propagated errors meaningless (Eqs. 4, 5). They are indicated in **Tables S2**, **S3** and **S4** to facilitate the comparison between phosphopeptides or between conditions, but should be used with caution.

#### Missing values and dissociation constants estimation

In several instances the phosphopeptide was quantified in the control but not quantified or quantified only once in the 14-3-3 replicates, preventing a *p*-value or even a FC value calculation and thus *K*_d_ determination. These missing values in the 14-3-3 replicates are due to low signals of unbound peptides, below the MS detection limit. Indeed, some well-known 14-3-3 phosphosites were in that case (see for examples the interactions between 14-3-3γ and pThr733 or pThr734 from RSK or pSer376 from SLP76 – phosphopeptides N°11, 12 and 19 in **Table S2**, respectively). As excluding those peptides based on their missing values would lead to exclude the highest-affinity binders, we devised the following procedure. If the phosphopeptide was measured 0 or 1 times in the 14-3-3 replicates, we imputed all missing values with a low conservative value of 10.000. Then, a *K*_d_ was simulated from the resulting intensities if they fulfilled the above-mentioned thresholds for p-value and FC. Finally, depending on the value of this simulated *K*_d_ we assigned the interaction to one of the following categories: *K*_d_ <1, <10 or <100 μM. In all other cases the interaction was classified as “non determined” (n.d.). If the phosphopeptide was measured twice in the 14-3-3 replicates and the interaction was classified as non-determined, we applied the same procedure. Please note that those are only upper limits of affinity; an interaction classified as “*K*_d_ <100 μM” could have a real *K*_d_ in the sub-micromolar range. With such procedure we estimated a total of 150 dissociation constants, including for the above-mentioned well known high-affinity binders.

#### Identification of outliers of the affinity trend between the different 14-3-3 isoforms

We identified potential outliers of the general affinity trend observed between the different 14-3-3 isoforms in two different ways. First, we calculated for each phosphopeptide a slope considering its ΔG for each 14-3-3 as *y* and the general 14-3-3 affinity classification (γ=1, β=2, η=3, ζ=4, τ=5, ε=6, σ=7) as *x* (**Table S3**). A mean slope and its associated standard error value was calculated over the 165 phosphopeptides for which a *K*_d_ was measured for the entire 14-3-3ome, and phosphopeptides which slope was below or above mean ± 3 SD (99.7% confidence) were considered as outliers. Then, to assess more precisely phosphopeptides that would be specific to 14-3-3σ, we assessed potential outliers of the distribution of the ΔΔG values calculated between γ and σ (mean ± 3 SD). Of note, the two procedures resulted in the identification of the same identical outliers (see Results).

### Benchmarking the Holdup Multiplex by competitive fluorescence polarization (FP)

In direct FP the fluorophore can interfere with affinity determination, especially when using small peptides. For that reason, we used competitive FP as an orthogonal method to benchmark our Holdup Multiplex measurements. By competitive FP we quantified the affinity of 16 phosphopeptides that we determined by Holdup Multiplex to bind to the 14-3-3s with affinities spanning 3 orders of magnitude (**Figure S7**), and of 6 peptides designed according to different consensus sequences, in the absence or in the presence of FCA (**Figure S2)**.

The detailed experimental procedure is described in^28^. Briefly, the affinity of the fluorescent tracer (fpB6 peptide, derived from the HSPB6 protein; gift of NN. Sluchanko) for each 14-3-3 protein was first measured in direct FP measurements using 8 increasing concentrations of 14-3-3, in the absence or in the presence of 100 μM FCA. Based on the direct titrations, the 14-3-3 concentration to be used in indirect FP was chosen to achieve a 80-100% complex formation, and this mixture was used for preparing a dilution series of the competitor (the unlabeled peptide under study; 8 different concentrations). Each fluorescent measurement was done in technical triplicates with a PHERAstar microplate reader by using 485 ± 20 nm and 528 ± 20 nm band-pass filters for excitation and emission, respectively. Analysis were carried out using ProFit, an in-house developed, Python-based fitting program that uses a Monte Carlo approach to take into account the experimental variability^50^. The dissociation constants of the direct and competitive FP experiments were obtained by first fitting the direct data with a quadratic binding equation then the competitive data with a competitive equation. The later uses several obtained parameters from the first fit, including the affinity of the labelled peptide for the 14-3-3^74^. The reported affinities and their standard deviations are calculated over 500–1000 independent fits of simulated datasets. All binding data and the obtained fits are provided in **Figures S2** and **S7**.

## Supporting information

Table S1 (Library design)

Table S2 (All the affinity measurements)

Table S3 (Comparison between the seven human 14-3-3)

Table S4 (Fusicoccin)

## List of Supplementary Material

**Figure S1.** The different 14-3-3s used in this study.

**Figure S2.** Competitive Fluorescent Polarization measurements of the interaction between 14-3-3γ and phosphopeptides designed according to the inferred sequence consensus.

**Figure S3.** Volcano plot ofHoldup Multiplex of the phosphopeptide library against 14-3-3 β, η, ζ, τ, ε and σ.

**Figure S4.** Frequency logos for all the seven human 14-3-3s.

**Figure S5.** Affinity-weighted frequency logos for all the seven human 14-3-3s.

**Figure S6.** Reproducibility of the Holdup Multiplex measurements.

**Figure S7.** Competitive Fluorescent Polarization measurements of the interactions between the different 14-3-3 proteins and the 16 phosphopeptides of the benchmark.

**Table S1** - Library design

**Table S2** - Normalized intensities from MS data and conversion to dissociation constants

**Table S3** - Binding comparisons of the seven human 14-3-3

**Table S4** - Effect of Fusicoccin-A on 14-3-3 / phosphomotifs interaction

## Acknowledgement

We are grateful to Goran Bich and Yves Nominé for their advises on with peptide design and data analysis, and to Nikolai N. Sluchanko for providing the FCA and the labeled fpB6 peptide. GG was supported by the *Post-doctorants en France* program of the *Fondation ARC* and by the collaborative post-doc grant of IGBMC. The Travé team was supported by the *Ligue Contre le Cancer* (équipe labellisée 2015 to GT) and the *Agence Nationale de la Recherche* (grant UBE3A ANR-18-CE92-0017 to GT). As a member of the IGBMC institute, the team also benefited from the French Infrastructure for Integrated Structural Biology (FRISBI) ANR-10-INSB-05-01, from Instruct-ERIC, from IdEx Unistra (ANR-10-IDEX-0002), from SFRI-STRAT’US project (ANR 20-SFRI-0012), and from EUR IMCBio (ANR-17-EURE-0023) under the framework of the French Investments for the Future Program as a member of the Interdisciplinary Thematic Institute IMCBio, as part of the ITI 2021-2028 program of the University of Strasbourg, CNRS and Inserm under the framework of the French Investments for the Future Program. The Carapito team is supported by the *Agence Nationale de la Recherche* (ANR-10-INBS-08-03) and the French Proteomic Infrastructure (ProFI FR2048).

## Author contributions

FD performed the MS sample preparation and measurements. 14-3-3 cloning was performed by CK and purification by AR and GG. PE synthetized some peptides. Holdup Multiplex and competitive fluorescent polarization experiments were performed by EM. FD, GG, CC and EM analyzed the results. GT conceived the original idea, and GT and CC obtained the funding. EM conceived and supervised the project. GG, CC, GT and EM wrote the paper. All authors reviewed the manuscript.

