## Supplementary Figures for "The Holdup Multiplex, an assay for high-throughput measurement of protein-ligand affinity constants using a mass-spectrometry readout"

**Figure S1.** The different 14-3-3s used in this study.

**Figure S2.** Competitive Fluorescent Polarization measurements of the interaction between 14-3-3γ and phosphopeptides designed according to the inferred sequence consensus.

**Figure S3.** Volcano plot ofHoldup Multiplex of the phosphopeptide library against 14-3-3 β, η, ζ, τ, ε and σ.

**Figure S4.** Frequency logos for all the seven human 14-3-3s.

**Figure S5.** Affinity-weighted frequency logos for all the seven human 14-3-3s.

**Figure S6.** Reproducibility of the Holdup Multiplex measurements.

**Figure S7.** Competitive Fluorescent Polarization measurements of the interactions between the different 14-3-3 proteins and the 16 phosphopeptides of the benchmark.

**A**

AviTag-His$_6$-MBP-TEVsite

AviTag-His$_6$-MBP-TEVsite-14-3-3 (x7)

MBP-14-3-3 (x7)

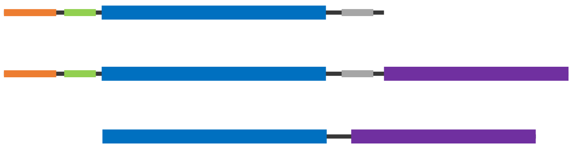

**B**

**AviTag-His$_6$-MBP-TEVsite**

MKGLNDIFEAQKIEWHEKHHHHHHPMKIEEGKLVIWINGDKGYNGLAEVGKKFEKDTGIKVTVEHPDKLEEKFPQVAATGDG
PDIIFWAHDRFGGYAQSGLLAEITPDKAFQDKLYPFTWDAVRYNGKLIAYPIAVEALSLIYNKDLLPNPPKTWEEIPALDKE
LKAKGKSALMFNLQEPYFTWPLIAADGGYAFKYENGKYDIKDVGVDNAGAKAGLTFLVDLIKNKHMNADTDYSIAEAAFNKG
ETAMTINGPWAWSNIDTSKVNYGVTVLPTFKGQPSKPFVGVLSAGINAASPNKELAKEFLENYLLTDEGLEAVNKDKPLGAV
ALKSYEEELAKDPRIAATMENAQKGEIMPNIPQMSAFWYAVRTAVINAASGRQTVDEALKDAQTNSSSNNNNNNNNNNPMSE
NLYFQGAMGSRG

**AviTag-His$_6$-MBP-TEVsite-14-3-3γ**

MKGLNDIFEAQKIEWHEKHHHHHHPMKIEEGKLVIWINGDKGYNGLAEVGKKFEKDTGIKVTVEHPDKLEEKFPQVAATGDG
PDIIFWAHDRFGGYAQSGLLAEITPDKAFQDKLYPFTWDAVRYNGKLIAYPIAVEALSLIYNKDLLPNPPKTWEEIPALDKE
LKAKGKSALMFNLQEPYFTWPLIAADGGYAFKYENGKYDIKDVGVDNAGAKAGLTFLVDLIKNKHMNADTDYSIAEAAFNKG
ETAMTINGPWAWSNIDTSKVNYGVTVLPTFKGQPSKPFVGVLSAGINAASPNKELAKEFLENYLLTDEGLEAVNKDKPLGAV
ALKSYEEELAKDPRIAATMENAQKGEIMPNIPQMSAFWYAVRTAVINAASGRQTVDEALKDAQTNSSSNNNNNNNNNNPMSE
NLYFQGAMVDREQLVQKARLAEQAERYDDMAAAMKNVTELNEPLSNEERNLLSVAYKNVVGARRSSWRVISSIEQKTSADGN
EKKIEMVRAYREKIEKELEAVCQDVLSLLDNYLIKNCSETQYESKVFYLKMKGDYYRYLAEVATGEKRATVVESSEKAYSEA
HEISKEHMQPTHPIRLGLALNYSVFYYEIQNAPEQACHLAKTAFDDAIAELDTLNEDSYKDSTLIMQLLRDNLTLWTSDQQD
DDGGEGNN

**MBP-14-3-3γ**

MKIEEGKLVIWINGDKGYNGLAEVGKKFEKDTGIKVTVEHPDKLEEKFPQVAATGDGPDIIFWAHDRFGGYAQSGLLAEITP
**AA**AFQDKLYPFTWDAVRYNGKLIAYPIAVEALSLIYNKDLLPNPPKTWEEIPALDKELKAKGKSALMFNLQEPYFTWPLIAA
DGGYAFKYENGKYDIKDVGVDNAGAKAGLTFLVDLIKNKHMNADTDYSIAEAAFNKGETAMTINGPWAWSNIDTS**A**VNYGVT
VLPTFKGQPSKPFVGVLSAGINAASPNKELAKEFLENYLLTDEGLEAVNKDKPLGAVALKSYEEELAKDPRIAATMENAQKG
EIMPNIPQMSAFWYAVRTAVINAASGRQTVD**A**AL**AA**AQTNAAAMVDREQLVQKARLAEQAERYDDMAAAMKNVTELNEPLSN
EERNLLSVAYKNVVGARRSSWRVISSIEQKTSADGNEKKIEMVRAYREKIEKELEAVCQDVLSLLDNYLIKNCSETQYESKV
FYLKMKGDYYRYLAEVATGEKRATVVESSEKAYSEAHEISKEHMQPTHPIRLGLALNYSVFYYEIQNAPEQACHLAKTAFDD
AIAELDTLNEDSYKDSTLIMQLLRDNLTLWTSDQQDDDGGEGNN

**Figure S1. The different 14-3-3s used in this study. (A)** Schematic representation of the different constructions used. **(B)** Sequence information. The sequences for the MBP control and 14-3-3γ constructs are indicated. The sequences of the other 14-3-3 constructs are obtained by substituting the 14-3-3γ sequence by their corresponding sequences from Uniprot. The MBP-14-3-3 constructs were initially optimized for crystallization and the MBP carries 5 Ala substitutions (in bold). All the sequences were validate by DNA sequencing at the plasmids level and by mass spectrometry at the purified proteins level. **(C; next page)** Sequence coverage achieved by LC-MS/MS peptide mapping on the 14-3-3 moiety for the different fusion proteins used.

**C**

| Protein | Application | 14-3-3 coverage |
|---|---|---|
| **14-3-3γ** | Holdup Multiplex | 89 % |
| **14-3-3γ** | competitive FP | 73 % |
| **14-3-3β** | Holdup Multiplex | 68 % |
| **14-3-3β** | competitive FP | 68 % |
| **14-3-3η** | Holdup Multiplex | 75 % |
| **14-3-3η** | competitive FP | 78 % |
| **14-3-3ζ** | Holdup Multiplex | 85 % |
| **14-3-3ζ** | competitive FP | 78 % |
| **14-3-3τ** | Holdup Multiplex | 78 % |
| **14-3-3τ** | competitive FP | 78 % |
| **14-3-3ε** | Holdup Multiplex | 93 % |
| **14-3-3ε** | competitive FP | 77 % |
| **14-3-3σ** | Holdup Multiplex | 86 % |
| **14-3-3σ** | competitive FP | 86 % |

**Figure S1 (continuing)**

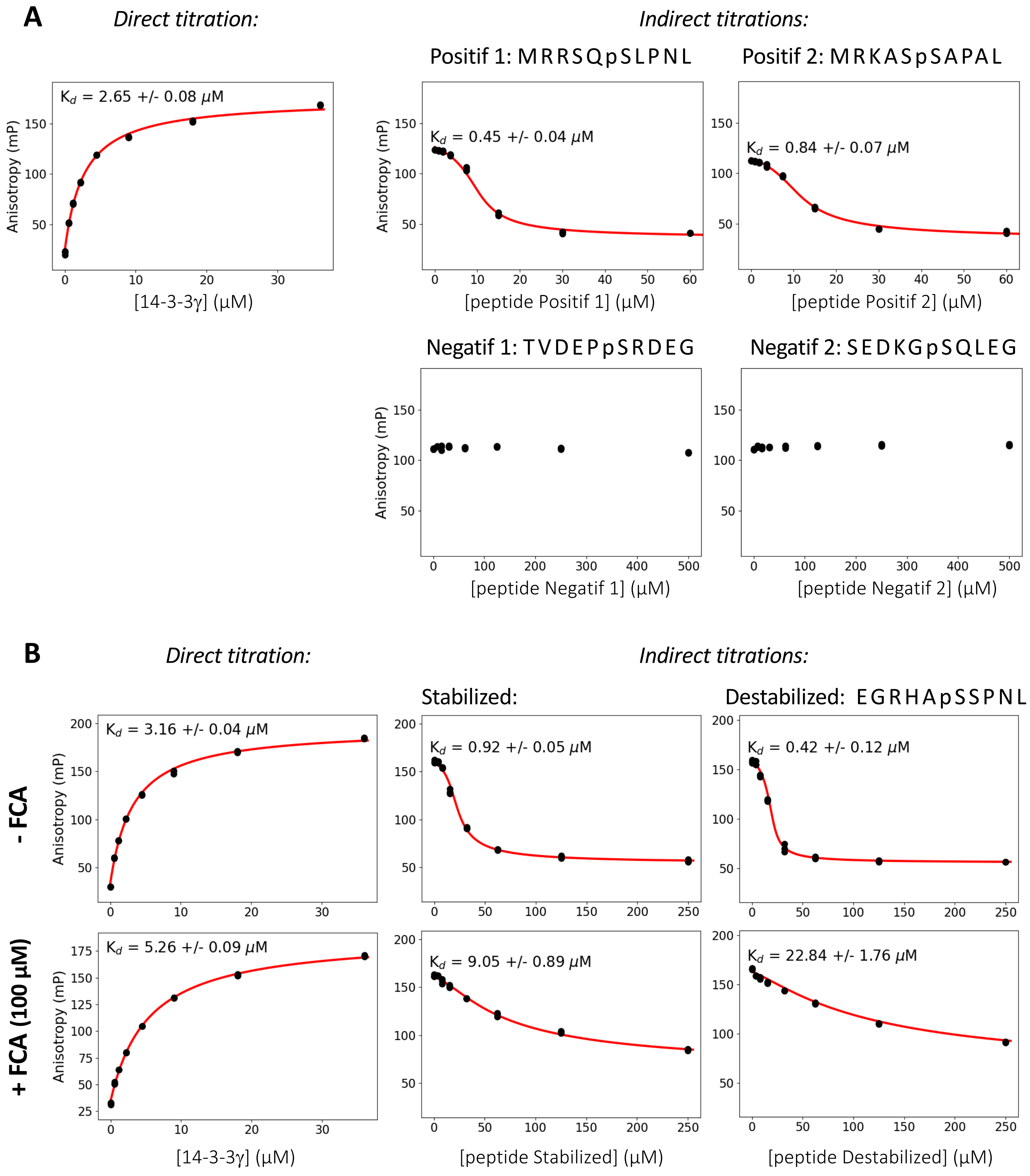


**Figure S2. Competitive Fluorescent Polarization measurements of the interaction between 14-3-3γ and phosphopeptides designed according to the inferred sequence consensus. (A)** Interaction with peptides designed according to the 14-3-3γ positive and negative consensus. Note the different peptide concentration ranges used for positive *vs* negative peptides. **(B)** Interaction with peptides designed according to the consensus for peptides which interaction with 14-3-3γ is stabilized or destabilized by FCA. Both direct and indirect titrations were performed with or without 100 μM FCA.

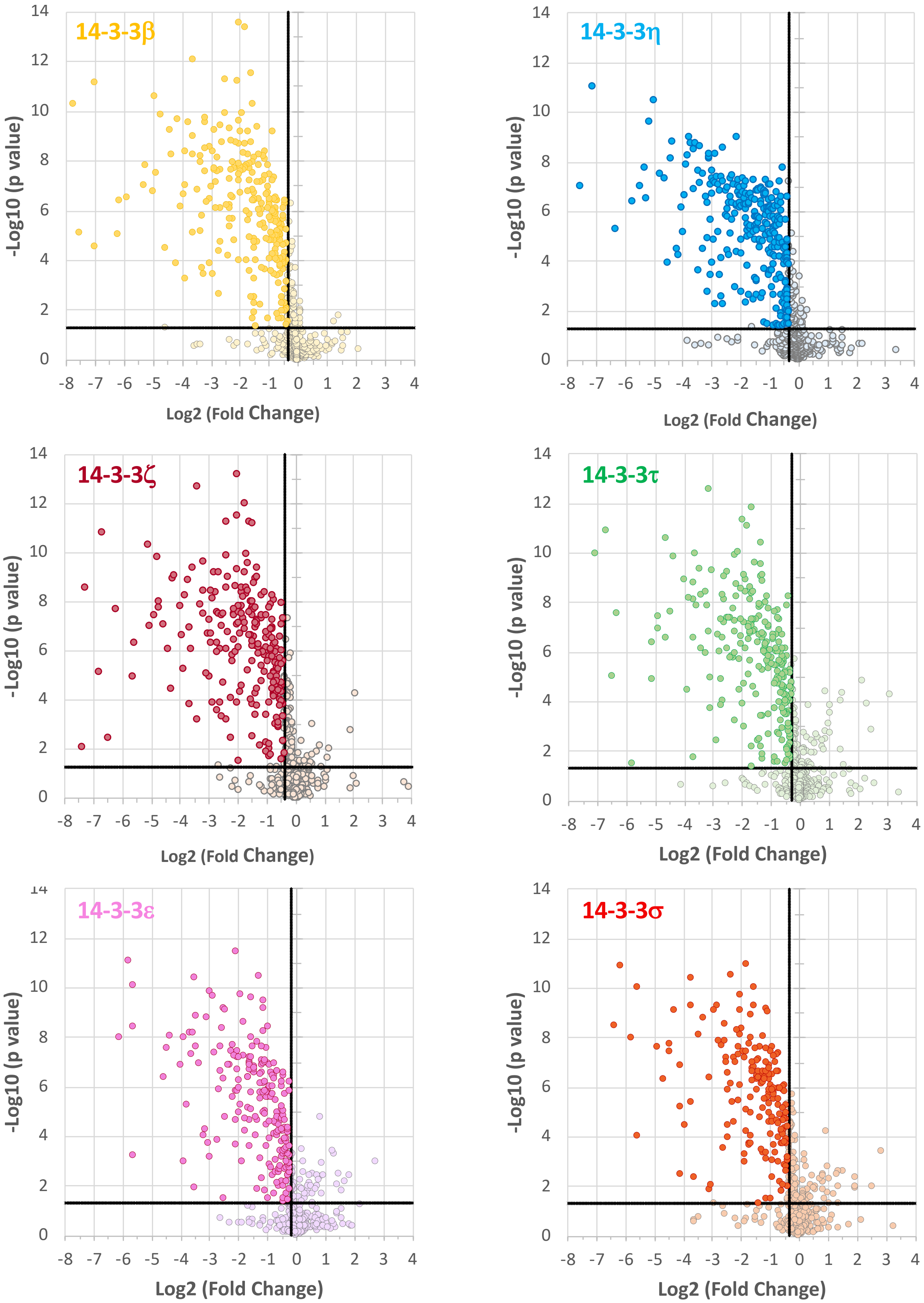


**Figure S3. Volcano plot of Holdup Multiplex of the phosphopeptide library against 14-3-3 β, η, ζ, τ, ε and σ.** The p-values (p-value ≤ 0.05) and fold-change ($Log_2(FC) \leq 2*SD\{Log_2(FC)\}$) thresholds used are indicated by plain lines. The fold-change thresholds change from one 14-3-3 to another, depending on the distribution of $Log_2(FC)$ (see Methods). Volcano plot for 14-3-3γ is presented Figure 2B.

**Figure S4. Frequency logos for all the seven human 14-3-3s**. Only the 563 phosphomotifs which interaction has been determined for all the 14-3-3 are considered. For each 14-3-3, the sequence logos highlight differences between phosphopeptides with p$K_d$ ≥ 3.50 *vs* p$K_d$ < 3.50; the number of phosphopeptides in each category is indicated within brackets.

**Figure S5 (next page). Affinity-weighted frequency logos for all the seven human 14-3-3s.** Only the 563 phosphomotifs which interaction has been determined for all the 14-3-3 are considered. The first panel represents the sequence logos of these 563 motifs. For each 14-3-3, affinity-weighted frequency logos are calculated over the N peptides (number in brackets) that have a quantified p$K_d$ ≥ 3.50. Contrarily to the sequences logos of Figure S4, here the estimated $K_d$ are not considered.

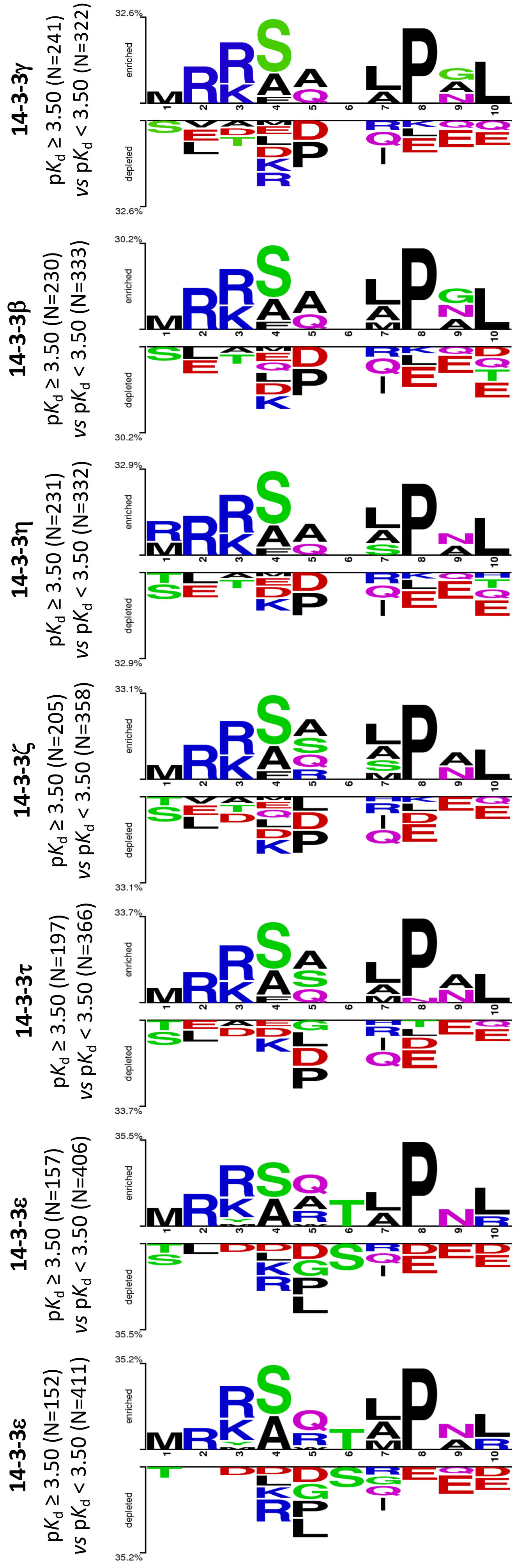

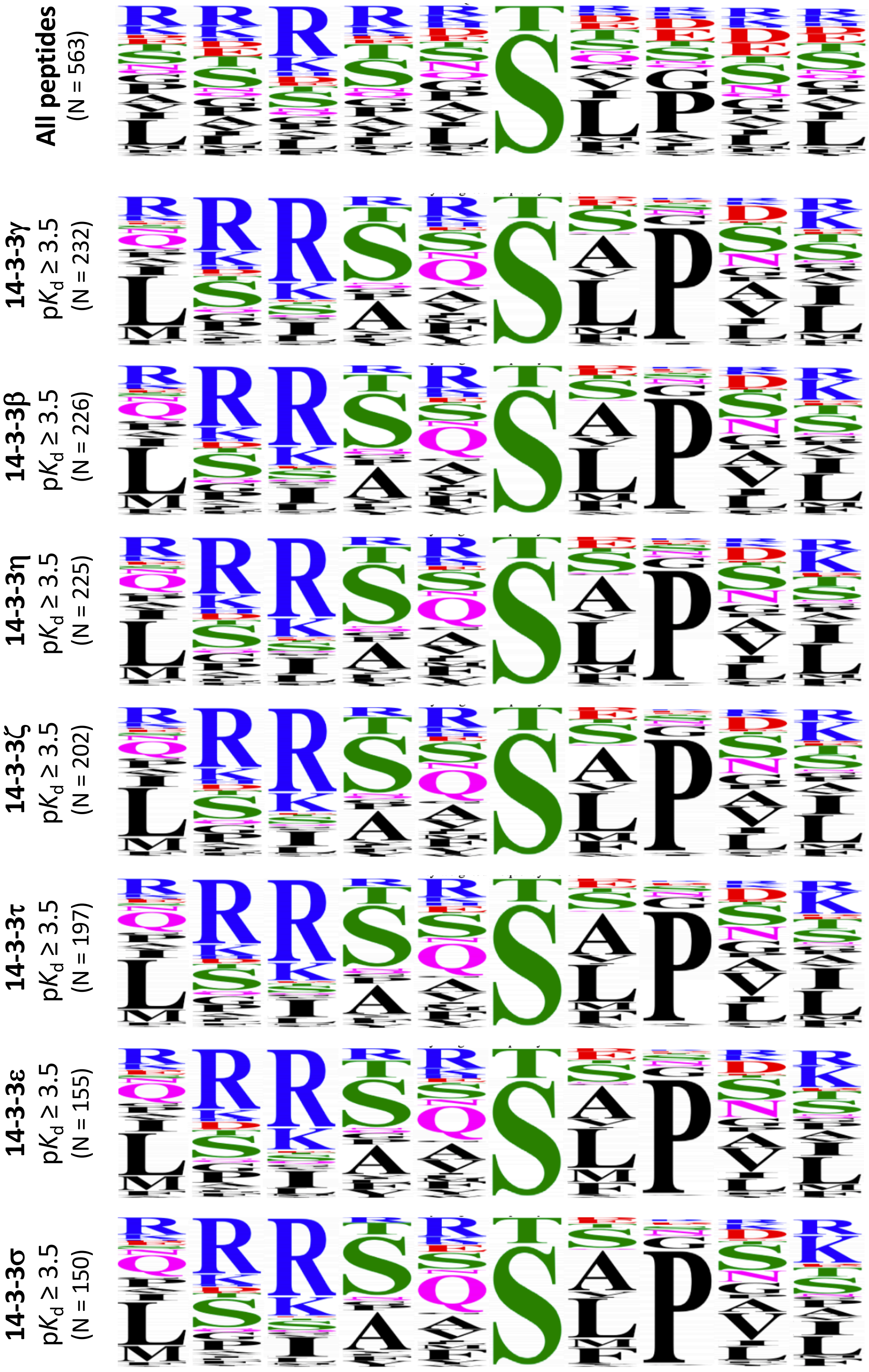


**Figure S5 (see legend in previous page)**

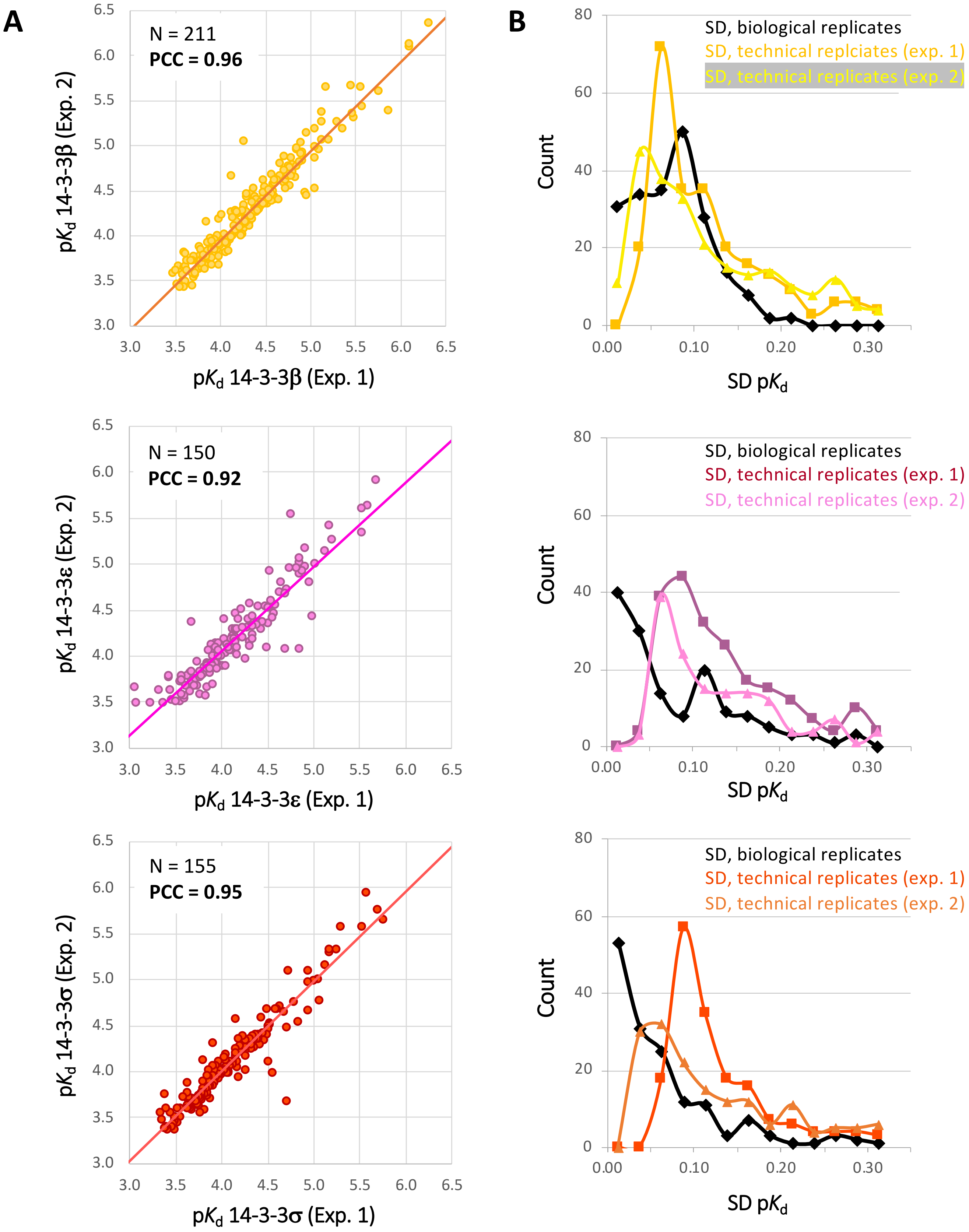


**Figure S6. Reproducibility of the Holdup Multiplex measurements. (A)** Correlations between the p$K_d$ measured in 2 independent experiments. **(B)** The p$K_d$ standard deviations between independent experiments (biological replicates) and inferred from one experiment (technical replicates) are similar. Results are shown for 14-3-3β, ε and σ; reproducibility for 14-3-3γ is indicated in Figure 5B.

**A**

**14-3-3γ**

Direct titration:

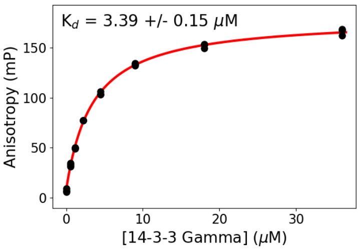


Indirect titrations:

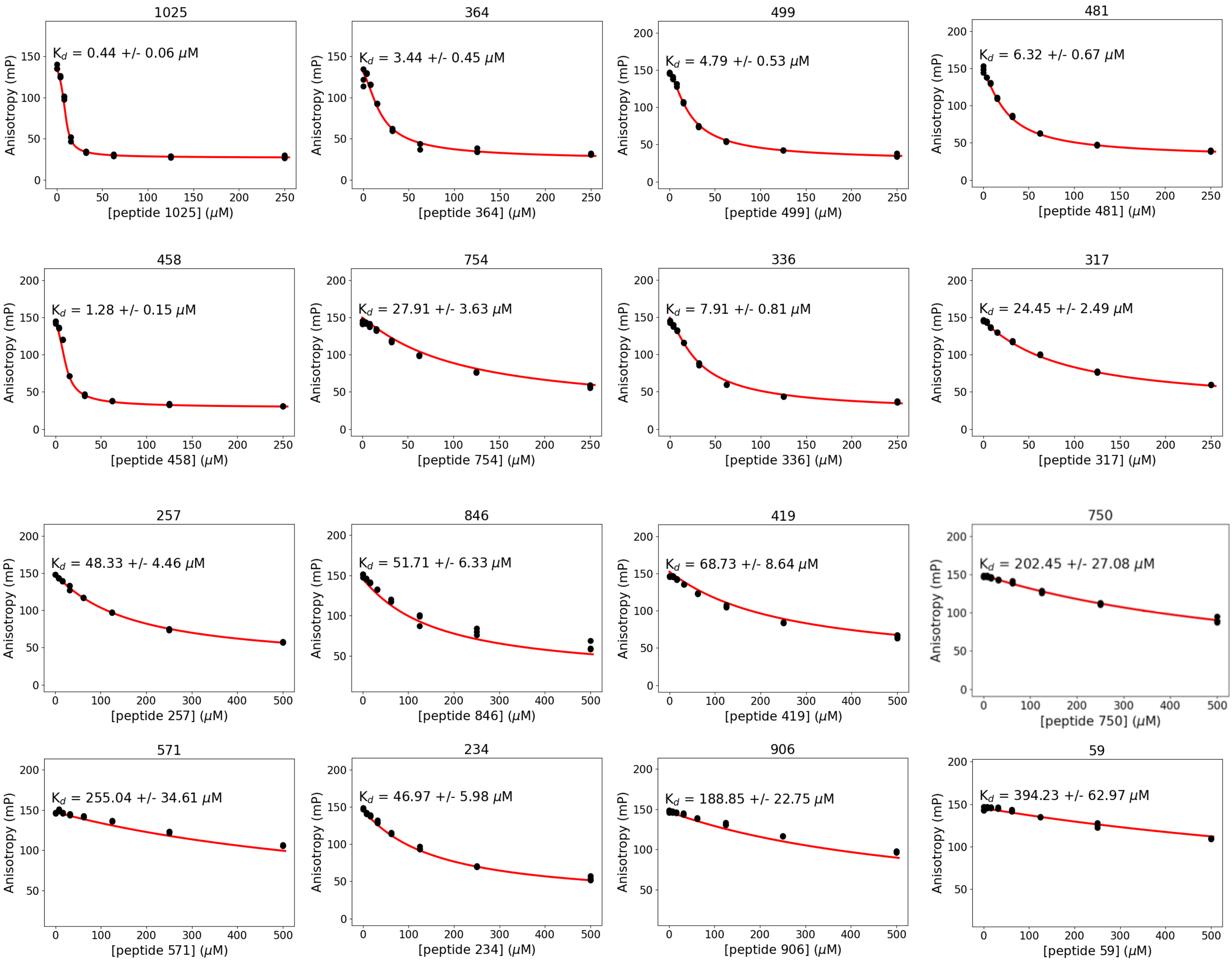


**Figure S7 (A to G; 7 pages). Competitive Fluorescent Polarization measurements of the interactions between the different 14-3-3 proteins and the 16 phosphopeptides of the benchmark.** For each 14-3-3, the upper right panel shows the direct FP experiment between the labeled peptide tracer and the titrated 14-3-3 protein and the following panels show competitive titrations. Competitive experiments were performed at a relatively high protein concentration to achieve 80% complex formation with the peptide tracer. Obtained polarization values were fitted with ProFit. No fitted curve is shown if we did not observe a quantifiable competition.

**B**

# 14-3-3β

Direct titration:

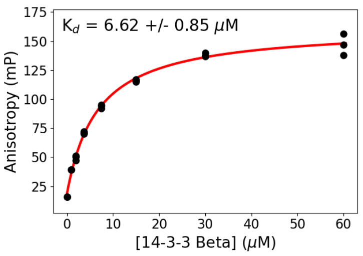


Indirect titrations:

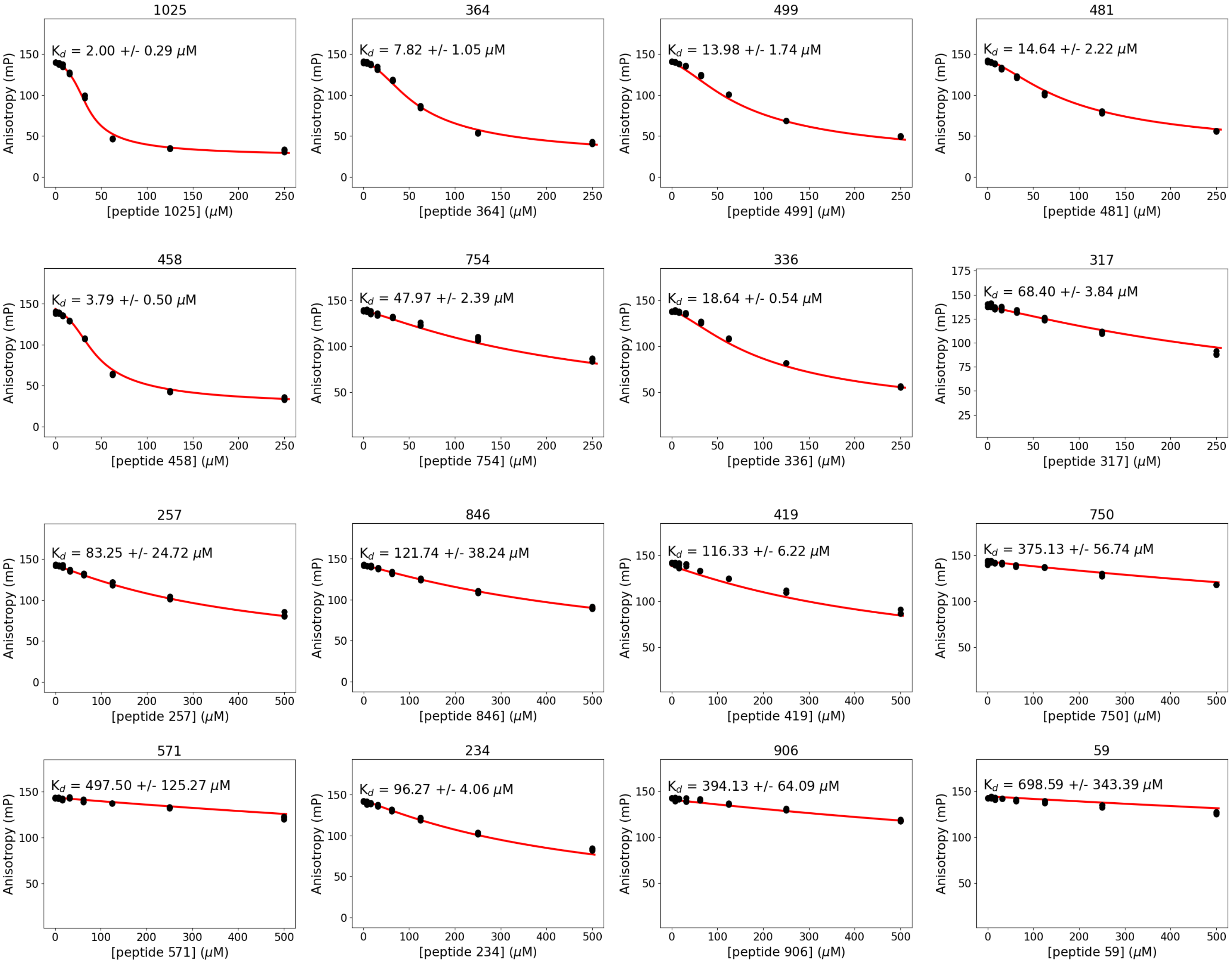


**Supplementary Figure 7.**

# C

## 14-3-3η

Direct titration:

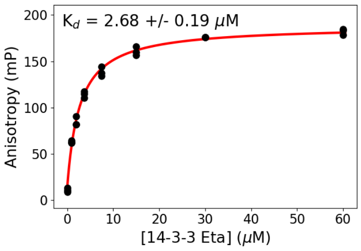


Indirect titrations:

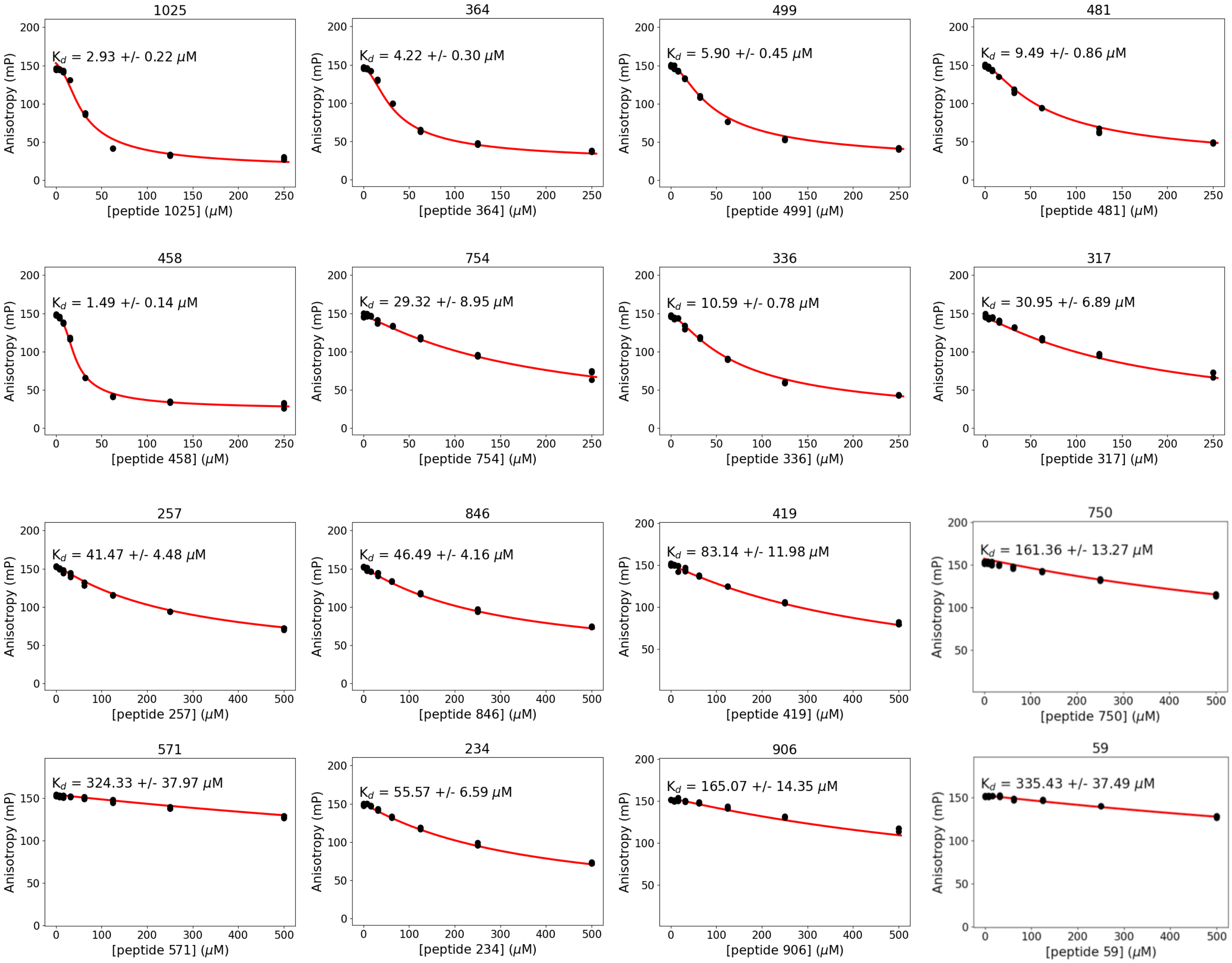


**Supplementary Figure 7.**

**D**

**14-3-3ζ**

Direct titration:

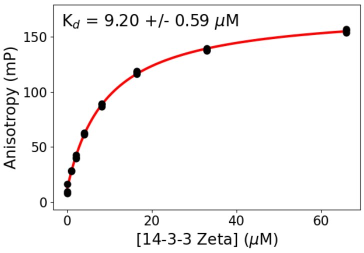


Indirect titrations:

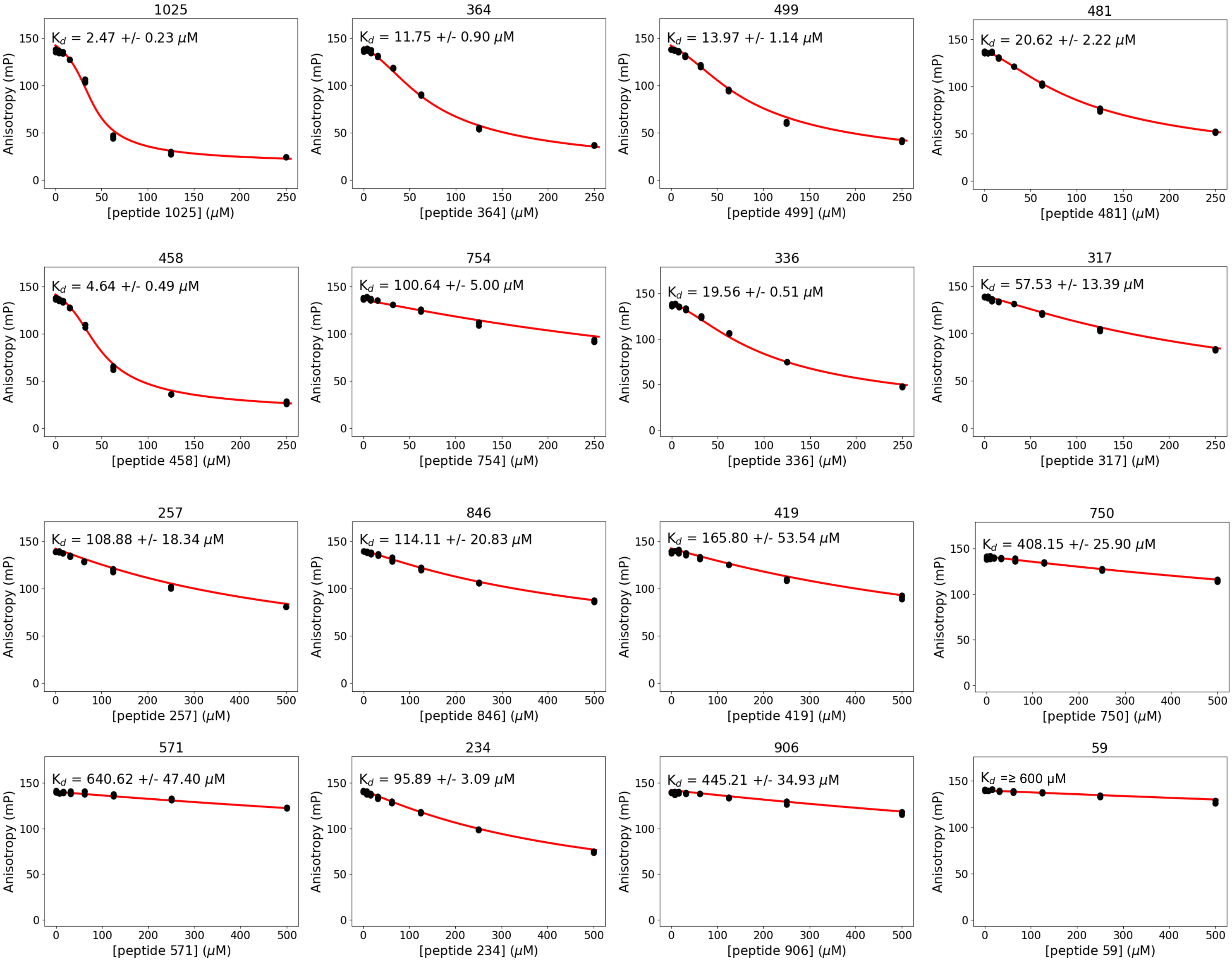


**Supplementary Figure 7.**

**E** **14-3-3τ**

Direct titration:

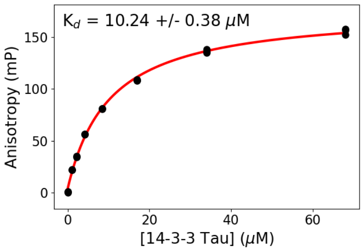


Indirect titrations:

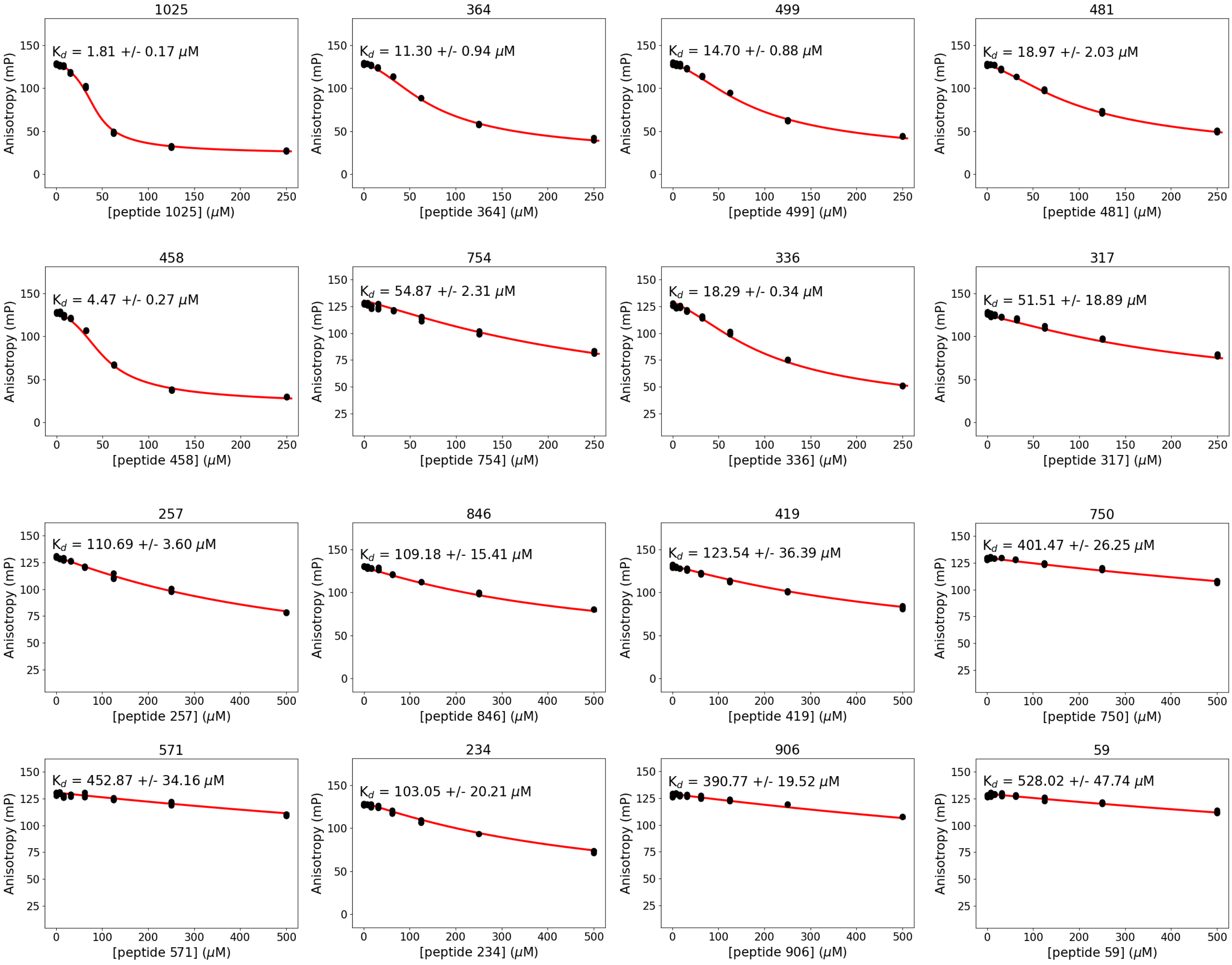


**Supplementary Figure 7.**

**F**

# 14-3-3ε

Direct titration:

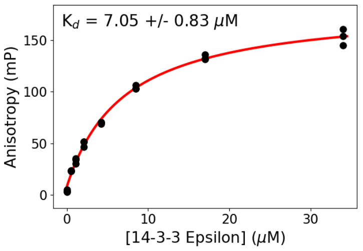


Indirect titrations:

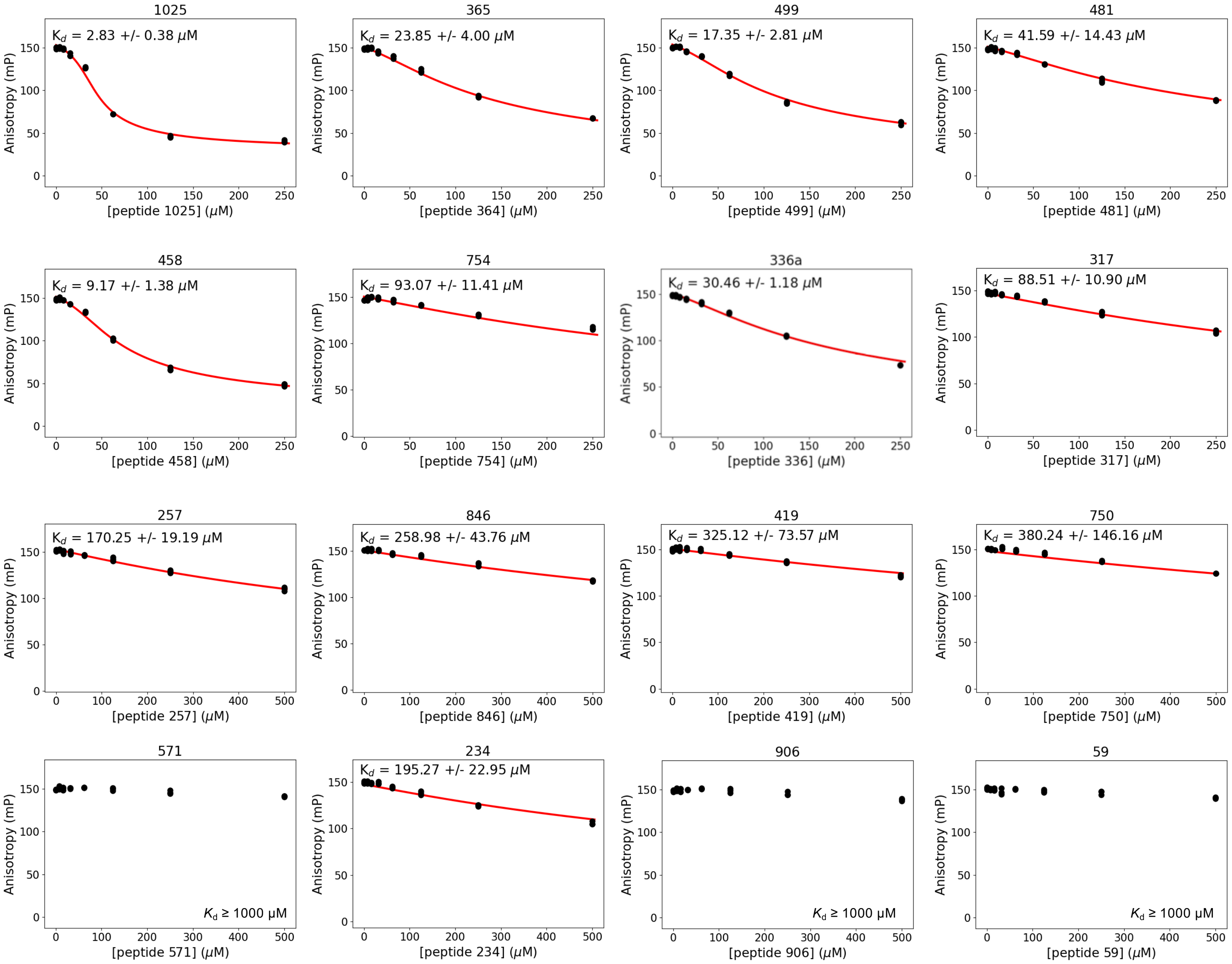


**Supplementary Figure 7.**

**G**

# 14-3-3σ

Direct titration:

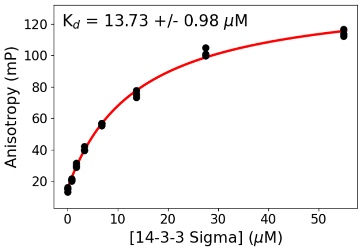


Indirect titrations:

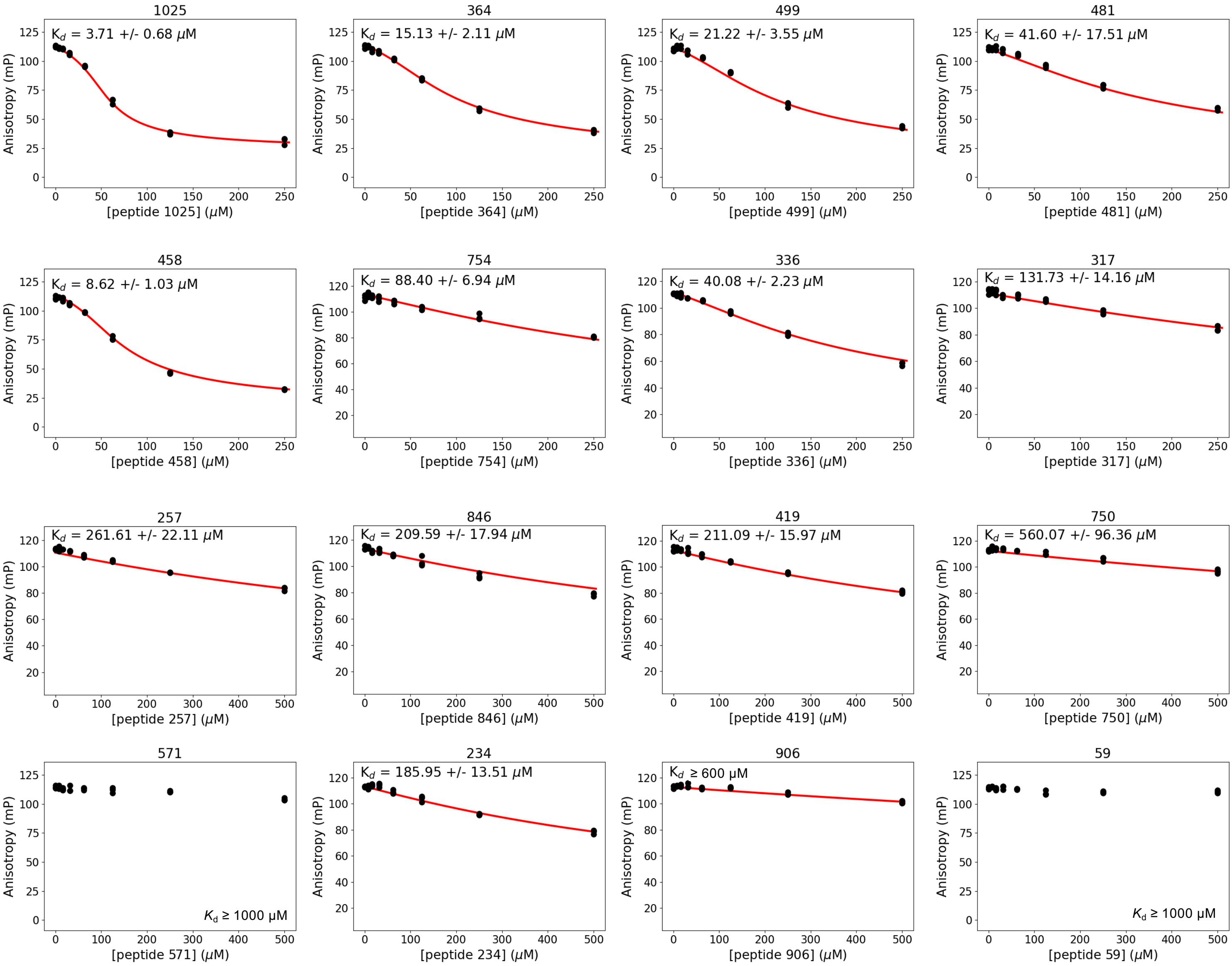


**Supplementary Figure 7.**
